# Contributions of Bayesian and Discriminative Models to Active Visual Perception across Saccades

**DOI:** 10.1101/2022.06.22.497244

**Authors:** Divya Subramanian, John Pearson, Marc A. Sommer

**Affiliations:** Department of Neurobiology, Duke School of Medicine, Duke University, Durham, NC, 27710, USA; Center for Cognitive Neuroscience, Duke University, Durham, NC, 27708, USA; Department of Biostatistics & Bioinformatics, Duke School of Medicine, Duke University, 27710, Durham, NC, USA; Department of Biomedical Engineering, Pratt School of Engineering, Duke University, Durham, NC, 27708, USA; Department of Psychology & Neuroscience, Trinity College of Arts and Sciences, Duke University, Durham, NC, 27708, USA

**Keywords:** vision, saccades, perception, active perception, Bayesian models, priors, discriminative models, primates, corollary discharge

## Abstract

The brain interprets sensory inputs to guide behavior, but behavior disrupts sensory inputs. In primates, saccadic eye movements displace visual images on the retina and yet the brain perceives visual stability, a process called active vision. We studied whether active vision is Bayesian. Humans and monkeys reported whether an image moved during saccades. We tested whether they used prior expectations to account for sensory uncertainty in a Bayesian manner. For continuous judgments, subjects were Bayesian. For categorical judgments, they were anti-Bayesian for uncertainty due to external, image noise but Bayesian for uncertainty due to internal, motor-driven noise. A discriminative learning model explained the anti-Bayesian effect. Therefore, active vision uses both Bayesian and discriminative models depending on task requirements (continuous vs. categorical) and the source of uncertainty (image noise vs. motor-driven noise), suggesting that active perceptual mechanisms are governed by the interaction of both models.

## Introduction

The process of generating percepts from external stimuli can be split into two stages. A stimulus is first detected by sensory receptors and encoded into neural signals^1^. Receptors provide evidence, *E*, for the stimulus, *S*, to the rest of the system. The evidence is then *decoded* to infer the stimulus from the evidence^2,3^. Under a probabilistic framework, the goal of decoding is to infer the probability of the stimulus given the evidence, *P*(*S*|*E*)^4^.

Models of decoding take two broad forms^5^. Discriminative models directly estimate P(S|E) by finding boundaries between evidence states and mapping stimulus states onto them^6,7^. Generative models, in contrast, build internal models of the world^8,9,10^. These include the joint probability of the stimulus and the evidence co-occurring, P(E, S). P(S|E) can be derived from the joint probability using Bayes’ rule. As such, Bayesian models are an implementation of generative models. Discriminative and Bayesian models have distinct strengths and are thought to combine for perception^11,12,13^. For example, discriminative models are flexible but vulnerable to sensory uncertainty. Bayesian models, on the other hand, use prior knowledge to optimally resolve uncertainty, e.g., due to environmental or receptor noise. Due to their robustness to noise, Bayesian models have been influential in explaining behavior across a wide range of sensorimotor stimulus estimation tasks^14–22^.

Sensory uncertainty may be introduced at the input stage or arise incidentally from one’s own movements. Constructing a stable, predictable percept of the world while moving through it constitutes *active perception*^23,24^. Active perception is fundamental for everyday behavior and its dysfunction may contribute to hallucinations and delusions^25,26,27^. Our overall goal was to test whether active perception, like many other systems, is explained by Bayesian models.

Vision across saccades is an apt model system for studying active perception^28,29^. Each saccade blurs and displaces the visual image on the retinas. To counter these disruptions, the visual system uses a copy of the saccade command, or “corollary discharge” to suppress the blur and nullify the displacements^30^. Previous work suggested that at least part of this process, saccadic suppression of the blur, is the outcome of combining motor and sensory information across saccades in a Bayes optimal manner^31,32^.

Here, we focus on whether Bayesian models are used to correct self-generated retinal shifts. The visual system, using corollary discharge, can remap its processing to predict its inputs after each saccade ^33,34,35^ and then compare that prediction with the postsaccadic visual input (Figure 1a). A match means that a viewed object was stable. Previous work by Rao et al.^36^ showed that humans use priors about the probability that an object will move for this process. They did not vary sensory uncertainty, however. Here we tested the hypothesis that the process of achieving perceptual stability across saccades is Bayesian. The key prediction is that priors about object movement are used more when sensory input is less certain.

**Figure 1.**
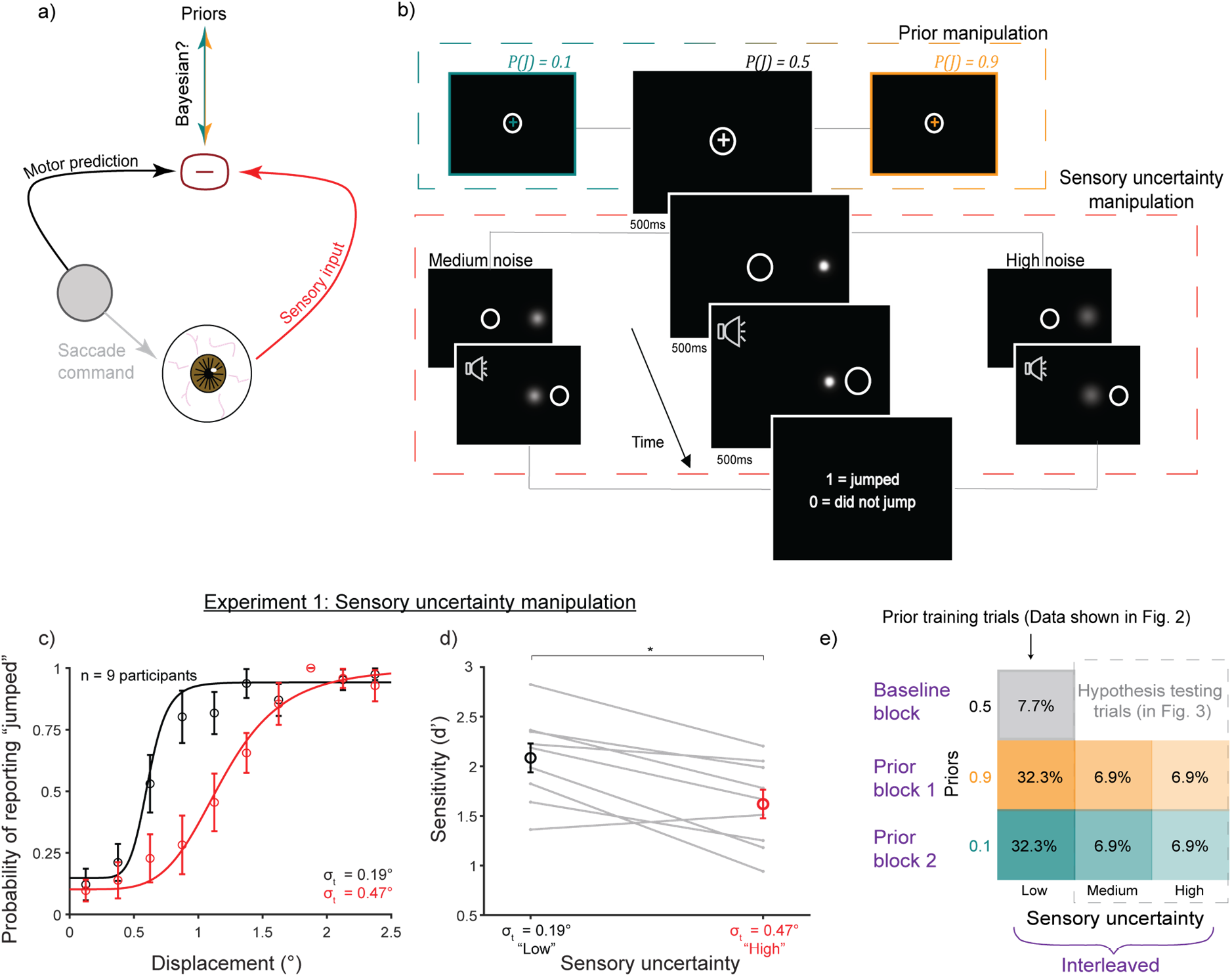
Experimental design. a) Judging whether an object is stable or moves during a saccade involves comparing a motor-driven prediction with sensory input. We tested if this process is Bayesian. b) Schematic of the SSD task. Middle, larger panels: “baseline” condition with neutral prior *P*(*J*)=0.5 and low uncertainty (minimal blur). White circle: eye position. High (0.9) and low (0.1) priors were cued by the color of the fixation cross (*top dashed box*). Sensory noise was manipulated by the width of the Gaussian target (*bottom dashed box*). c-d) Experiment 1 results (n=9 participants). c) Psychometric curves flattened and d) sensitivity decreased with greater Gaussian width σ_t_. *, p < 0.0125. e) Trial breakdown for Experiment 2. Blocks of high and low priors followed a baseline block. 70% of trials in the prior blocks were prior-training trials with low uncertainty and priors matched to true jump probability (results shown in Figure 2). 30% were hypothesis testing trials with medium and high uncertainty targets. Fixation colors cued the learned priors even though the true jump probability was 0.5 (results shown in Figure 3).

## Results

### Categorical judgments of displacement are anti-Bayesian

We used a modified Saccadic Suppression of Displacement (SSD) task^37^. Human participants fixated near the center of a screen, and upon being cued, made a saccade to a target. During the saccade, the target was displaced by varying amounts. After the saccade, participants reported their perception of whether the target had moved or not. The target always moved, but the displacement was drawn from two different distributions. On “jump trials,” it was drawn from a broad Gaussian distribution (μ=0°, σ=1.5°), and on “non-jump” trials it was drawn from a very narrow Gaussian distribution (μ=0°, σ=0.017°) (Figure S1). Since the distributions overlapped, there was no objectively correct solution to the task. The optimal solution would be to learn the relative probabilities of jump and non-jump trials. Information about the probability of jump trials was the “prior” for this task. To train priors or cue learned priors, we used the color of the fixation cross (Figure 1b, top dashed box), similar to the method of Rao et al.^36^

In Experiment 1 (n=9 participants), we first identified a stimulus manipulation that reliably induced sensory uncertainty (Figure 1b, lower dashed box). We tested four potential manipulations at two noise levels each and found that sensory uncertainty was most reliably induced by increasing the standard deviation of an isoluminant Gaussian “blob” target (Figure 1c,d). This yielded psychometric functions (fit to pooled data across participants; individual curves in Figure S2) that were steeper in the lower-noise condition (Figure 1c, black curve, σ_t_=0.19°) than in the high-noise condition (Figure 1c, red curve, σ_t_=0.47°). There were significant differences in d’ (ref. 38) between the low (mean=2.08, SE=0.15) and high (mean=1.62, SE=0.14) noise levels (p=0.0023; paired t-tests; α=0.0125 to correct for four comparisons) (Figure 1d). Figure S3 shows results for the other stimuli.

In Experiment 2, participants were trained on priors, *P*(*J*), using performance-based feedback. They first completed a baseline block of 100 trials in which the fixation cross was white and *P*(*J*) = 0.5. Two “prior blocks” of 600 trials, subdivided into training and testing trials, followed. Prior training trials constituted 70% of trials in the block (Figure 1e). The fixation color indicated *P(J*) (= 0.1 or 0.9) and the target had the lowest uncertainty (σ_t_=0.1°), essentially punctate. The remaining 30% of trials were “hypothesis testing” trials (Figure 1c). The color of the fixation cross indicated the prior, but the true jump probability was a neutral 0.5 to isolate the effects of the learned, color-cued priors. Targets in these trials had additional sensory uncertainty (“medium” or “high”) corresponding to wider Gaussians (σ_t_=0.25° and 0.5°, respectively). The hypothesis testing trials were relatively infrequent and interspersed randomly to mitigate the possibility of participants recognizing that higher noise targets implied a neutral prior. We also performed a control experiment in which the jump probability matched the priors across noise conditions (results in Figure 3).

To analyze data from Experiment 2, we compared participants’ performance with the predictions of a Bayesian ideal observer model (Figure 2a-c; details under *Detailed Methods: Section 4*). On every trial, the ideal observer decided whether the probe jumped or not given a perceived displacement, 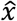. The decision would be “yes” if the probability of a jump given 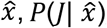, exceeded the probability of non-jump given 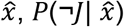:

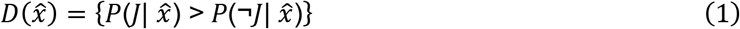

**Figure 2.**
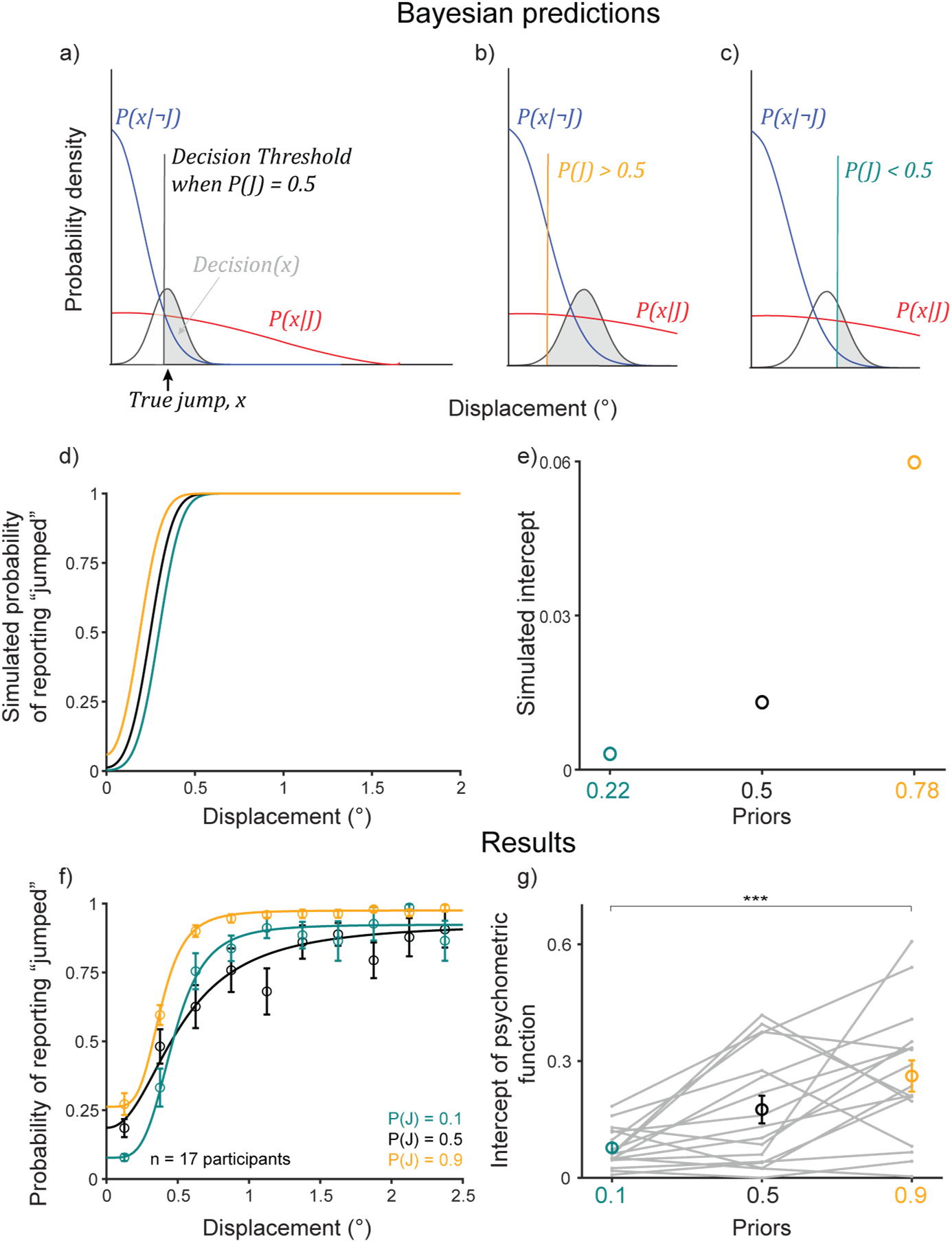
Participants learned the priors. a-c) Bayesian ideal observer models. d) Bayesian predictions for prior learning. e) Intercepts for curves in (d). f) Psychometric curves from n=17 participants. Bins averaged across participants. Error bars: S.E.M. Curves were fit to pooled data. g) Intercepts for curves in (f), fit to individual participants, matched Bayesian predictions in (e). Gray lines: individual participants. Colored markers and error bars: means and SEMs across participants. ***, p < .001.

The decision variable 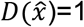 (jumped) if the condition in braces is met. Otherwise, 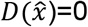 (no jump). Using Bayes’ rule,

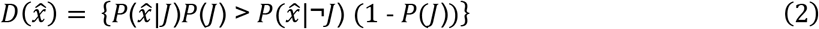

where 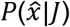 and 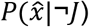 are the likelihoods of “jump” and “no jump,” respectively. For each prior, *P*(*J*), there was a threshold at which the condition in braces was met, i.e., it was equally likely that the probe jumped or did not. For *P*(*J*)=0.5, it was where the two likelihood distributions intersected and were equal (Figure 2a, black vertical line). If there was no sensory uncertainty, i.e., if 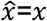 where *x* was the true displacement, the ideal observer would report “no jump” for all displacements less than the threshold and “jump” for all displacements above the threshold. Since the target was a Gaussian blob, we assume Gaussian uncertainty *σ_t_*, determined by the target, about the true displacement, *x* (Figure 2a, black distribution):

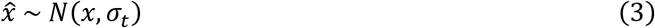

Therefore, the decision given the true displacement, *D*(*x*), was the integral over values of *x* greater than the decision threshold (Figure 2a, shaded region):

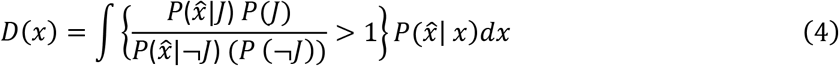

This restricted the value of the decision to range from 0 to 1.

We first assessed prior learning. For a high prior, e.g., *P*(*J*)=0.9, the threshold would move to the left since 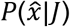 was weighted higher than 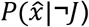, thus increasing the ratio in the braces (Figure 2b), and vice versa for a lower prior, e.g. *P*(*J*)=0.1 (Figure 2c). Critically, for the same perceived displacement, the ideal observer was more likely to report that the probe jumped for a higher prior than for a lower prior. Figure 2d shows simulations for an ideal observer with likelihood distributions, 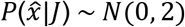 and 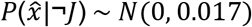, prior *P*(*J*)=0.22 (teal), *P*(*J*)=0.5 (black), and *P*(*J*)=0.78 (orange), and sensory noise, 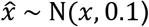. We chose *P*(*J*)=0.22 and 0.78 for the simulations to account for 70% true-statistic trials and 30% neutral. The key point was that the high prior was >0.5 and the low prior was <0.5. Figure 2e shows the value of the curves at displacement = 0 (the intercept).

For the human participants (n=17, Figure 2f), in the prior *training* trials, psychometric curves shifted upward for the high prior (orange curve) and downward for the low prior (teal curve) at small displacements as predicted. The crossing of the low-prior (teal) curve over the black was not predicted but has implications that are addressed later (Figure. 8 and its associated text). Data for individual participants are shown in Figure S4. A lower intercept in the low prior condition than in the high prior condition (Figure 2g) matched the Bayesian predictions in Figure 2e. Repeated-measures ANOVA on the intercepts with prior as the within-conditions factor yielded a significant main effect of priors (F(2)=11.82; p=0.0001). Post-hoc comparison (Tukey HSD) of the *P*(*J*)=0.9 and 0.1 conditions, the two priors tested later in hypothesis testing trials, showed that high-prior intercepts (mean=0.26, SE=0.04) were significantly higher than low-prior intercepts (mean=0.08, SE=0.01; p=2.79×10^−4^). These results indicated that participants learned the priors as expected.

In the randomized, less frequent *hypothesis testing* trials, we tested the Bayesian hypothesis that priors are used more with increasing uncertainty. In these trials, the targets had medium or high sensory uncertainty. Figures 3a,b show Bayesian predictions for these medium- and high-noise conditions, respectively. We used the same likelihood ratios and priors as in Figure 2a-c, but with sensory noise 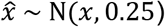 and 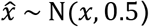, respectively, to match the medium and high noise target widths. The model predicted greater separation between the low- (teal) and high-prior (orange) decision curves, i.e., greater prior use, in the high noise condition than in the medium noise condition, quantified by the high prior - low prior intercept difference (Figure 3c). In other words, the Bayesian ideal observer used the prior more with increasing sensory noise.

**Figure 3.**
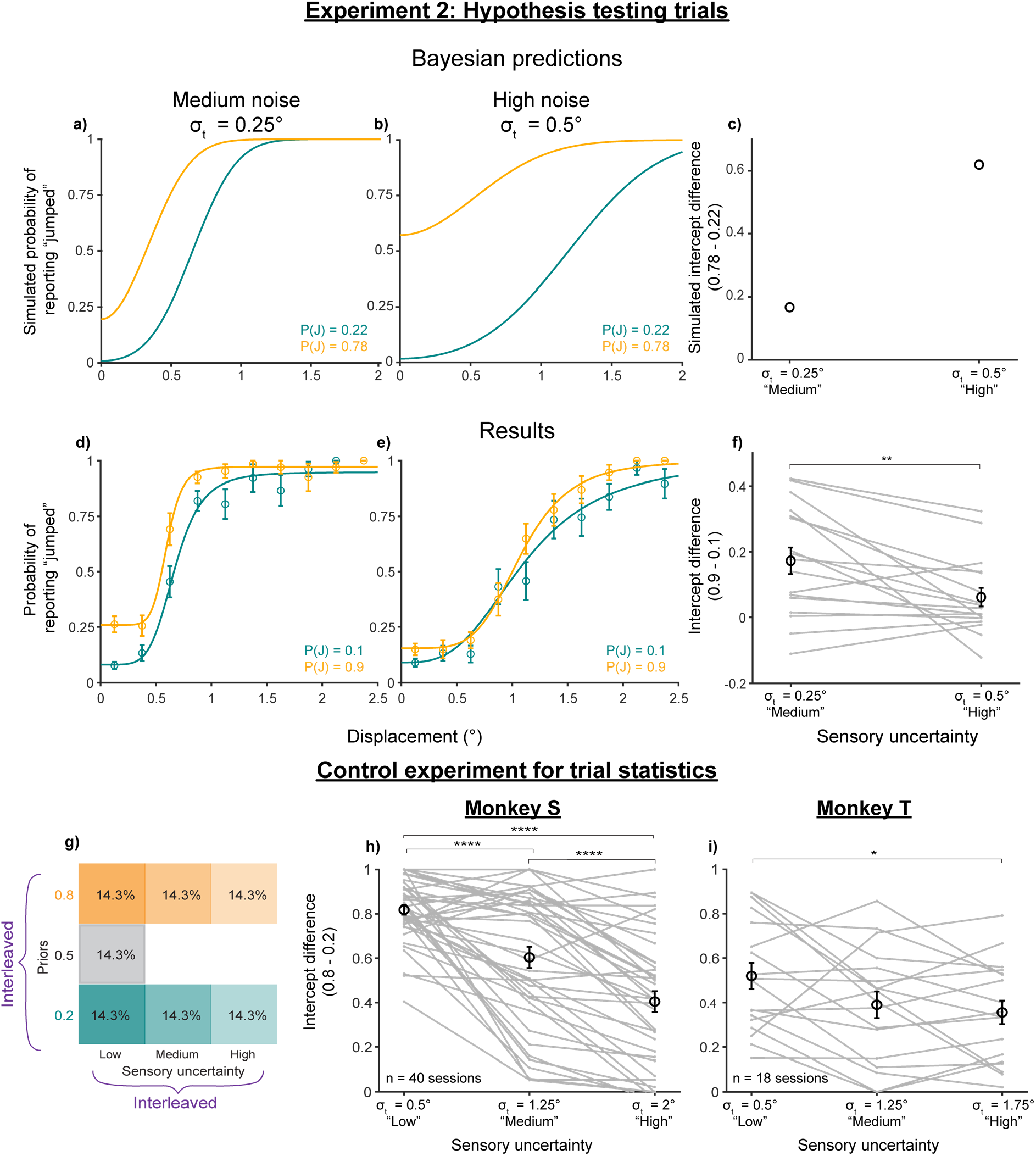
Categorical judgments of displacement are anti-Bayesian. a-b) Predicted psychometric curves from the Bayesian ideal observer model for the (a) medium and (b) high noise conditions. c) High-low prior intercept differences for the curves in a-b. d-f) Results from n=17 participants for the (d) medium and (e) high noise conditions, and (f) the respective high-low prior intercept differences. **, p<0.01. g-h) Results from a control experiment run on monkeys, in which the true jump probability matched the prior for the medium- and high-noise trials. g) Trial breakdown. h-i) High-low prior intercept difference) across noise levels for (h) Monkey S and (i) Monkey T. ****, p<0.0001; *, p<0.05.

Human participants showed the opposite effect: they used their priors *less* with increasing noise. Psychometric curves across priors moved closer together in the high-noise (σ=0.5°) condition (Figure 3e) compared to the medium-noise (σ=0.25°) condition (Figure 3d). Data for individuals shown in Figures S5 and S6. The difference in intercepts was significantly greater in the medium-noise condition (mean=0.17, SE=0.04) than in the high-noise condition (mean=0.06, SE=0.03; p=0.0081 using a paired t-test) (Figure 3f). Overall, the results in Experiment 2 suggested that, in this sense, human participants were *anti-Bayesian*.

We considered the possibility that participants were not anti-Bayesian but had learned that trials with medium- and high-noise targets had a neutral jump probability. In this case, their prior for the hypothesis testing trials would be 0.5 and the Bayesian prediction is for the orange and teal psychometric curves to collapse together with increasing noise. Note that if the participants *only* learned the priors according to target type (i.e., low noise targets = color-cued prior, but medium and high noise targets = 0.5), then there would be no separation between the orange and teal psychometric curves at all. Therefore, participants clearly learned the color-associated priors. Nevertheless, to account for this potential confound, we analyzed results from a control experiment in two rhesus macaques in which the jump probability matched the color-associated prior for *all* noise levels. A full description of the monkey experiments is provided in *Detailed Methods: Section 2*. Briefly, all seven trial types (3 priors with low sensory noise + 2 each with medium and high noise) were randomly interleaved and had the same relative frequencies (Figure 3g). Consistent with the human results, the intercept differences between the *P*(*J*)=0.8 and 0.2 conditions *decreased* with increasing sensory noise (3h-i) for both monkeys. Repeated-measures ANOVA on intercept differences with noise levels as the main within-subjects factor yielded significant effects (Monkey S: F(2)=51.75, p=4.97×10^−15^; Monkey T: F(2)=4.56, p=0.0176). For monkey S (n=40 sessions), post-hoc comparisons (Tukey HSD) showed that intercept differences in the low noise condition (σ_t_=0.5°; mean=0.82, SE=0.02) were significantly higher than in the medium noise (σ_t_=1.25°; mean=0.60, SE=0.05; p=3.49×10^−6^) and high noise (σ_t_=2°; mean=0.40, SE=0.05; p=0) conditions. Intercept differences in the medium noise condition were also higher than in the high noise condition (p=1.51×10^−5^). For Monkey T (n=18 sessions), there was a significant difference between the low (σ_t_=0.5°; mean=0.52, SE=0.06) and high noise (σ_t_=1.75°; mean=0.36, SE=0.05; p=0.0190) conditions. Intercept differences in the medium noise condition (σ_t_=1.25°; mean=0.39, SE=0.06) fell between the low and high noise conditions, not significantly different from either. Overall, the results replicated our human findings to confirm that the anti-Bayesian effect was based on learned, color-associated priors.

### Continuous judgments of displacement are Bayesian

Do the above results mean that the perception of visual displacement across saccades is always anti-Bayesian? Or was that outcome due, at least in part, to the categorical (binary) nature of the task? We tested this in Experiment 3 by requiring *continuous* estimates of displacement across saccades^31^. Human participants performed the same SSD task, but instead of providing a binary report of “jumped” or “did not jump,” they provided a continuous report using a mouse cursor (Figure 4a). The target jumps were horizontal, and the mouse cursor was restricted to that dimension. Formulating the task as a unidimensional, continuous problem allowed us to cast it in a form that has been tested across many sensorimotor domains^14,15,18,21,22^. If the uncertainty about the stimulus is modeled as the sensory likelihood, then the mean of the posterior (its maximum value and thus, our approximation of the inferred response) would be a reliability-weighted combination of the sensory likelihood and prior distributions (Figure 4b):

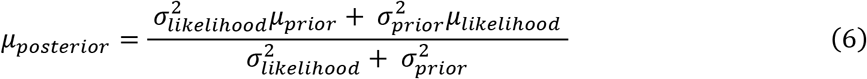

**Figure 4.**
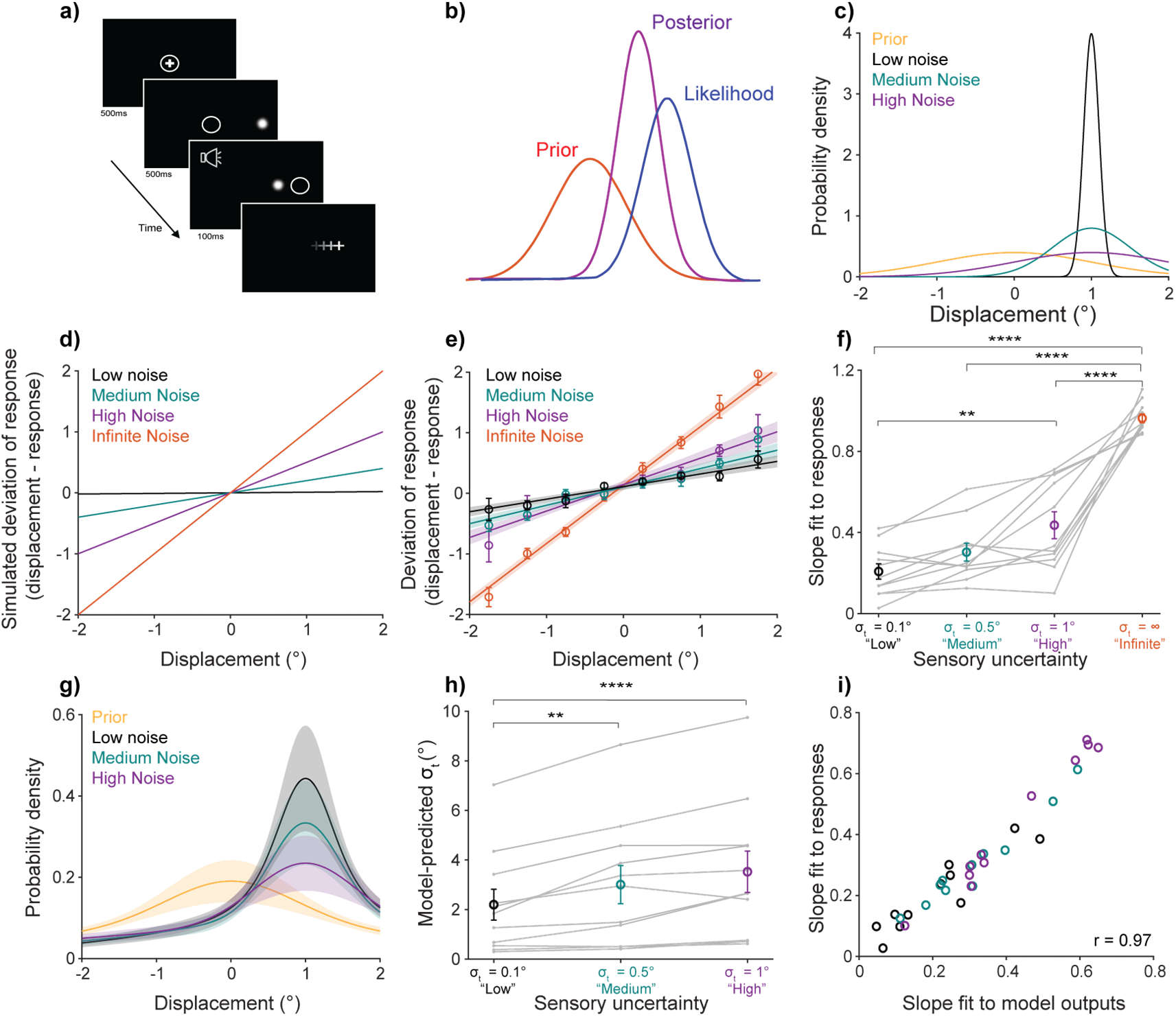
Continuous displacement perception is Bayesian. a) Task schematic. Participants performed the same SSD task as in Experiments 1 and 2 but provided a continuous estimate of where the target landed after the saccade using a mouse cursor (+). b) Bayesian inference for continuous stimulus estimation. c) Distributions used in the experiment. d) Bayesian predictions for the experimental parameters in (c). e) Results from n=11 participants. Bins and lines, fit to individuals, were averaged across participants. Error bars and shaded regions: S.E.M. f) Slopes of lines in (e). g) Model fits for participants’ internal prior and likelihood distributions, to be compared with (c). h) Standard deviations of likelihood distributions in (g). i) Correlation between model-predicted slopes computed in the same way as the slopes in (f) and the empirical slopes from (f) on a participant-by-participant basis. **, p<0.01; ****, p<0.0001.

As σ^2^_likelihood_ increases, with the other terms held constant, μ_posterior_ approaches μ_prior_. In other words, for a given prior with fixed uncertainty, the response should get closer to the prior with greater sensory uncertainty.

The prior was a Gaussian statistical distribution with μ_prior_=0° and σ_prior_=1°. Participants were first trained on the prior for 600 trials using performance-based feedback. They then performed 400 hypothesis testing trials that provided no feedback. There were four sensory uncertainty conditions: low noise (σ_t_=0.1°), medium noise (σ_t_=0.5°), high noise (σ_t_=1°), and an “infinite noise” condition in which the target did not reappear postsaccadically. Figure 4c illustrates the distributions used in the experiment, with the prior centered at 0° and the likelihood (Gaussian blob) distributions centered, for purpose of illustration, on displacement = 1°.

Figure 4d shows the predicted deviation in response from the presented displacement (displacement - response) for a Bayesian ideal observer (details in *Detailed Methods: Continuous Bayesian simulations*). If the sensory uncertainty is much smaller than that of the prior, as in the lowest noise condition (black line), then the deviation of the posterior (response) from the true displacement should be near 0 for all presented displacements. Conversely, for maximal sensory uncertainty as in the infinite noise condition (orange line), the response should always be the mean of the prior. Since μ_prior_=0°, the deviation for each displacement equals the displacement itself. The medium (teal) and high (purple) noise conditions are predicted to fall in between the low and infinite noise conditions, with slopes proportional to noise level. In summary, the slope of the deviation line increases with increasing sensory uncertainty.

Participants’ (n=11) responses in the hypothesis testing trials matched Bayesian predictions (Figure 4e). The slopes of the deviation lines increased with increasing sensory uncertainty (Figure 4f). Repeated-measures ANOVA on the slopes yielded a significant main effect of noise level (F(3)=78.01, p=2.87×10^−11^). Post-hoc comparison of conditions (Tukey HSD) showed that slopes in the low-noise (σ_t_=0.1°) condition (mean=0.21, SE=0.04) were significantly lower than in the high-noise (σ_t_=1°; mean=0.44, SE=0.07, p=0.0011) and infinite noise conditions (mean=0.96, SE=0.02; p=6.98×10^−14^). Also, slopes in the medium noise condition (σ_t_=0.5°, mean=0.30, SE=0.04) were significantly lower than in the infinite noise condition (p=2.05×10^−12^), and slopes in the high-noise condition were significantly lower than in the infinite-noise condition (p=4.65×10^−10^).

We fit individual participant responses to a Bayesian ideal observer model by minimizing squared error to infer their used prior and sensory likelihood distributions (Figure 4g). The prior mean and standard deviation, and standard deviations of the low-, medium- and high-noise parameters, were fit simultaneously by assuming that the response in the infinite noise condition was the prior mean. Model outputs for the prior mean (mean=0.05, SE=0.04) were not significantly different from 0, the true prior (p=0.22, one-sample t-test). Fit parameters for likelihood standard deviations increased with increasing noise, with repeated-measures ANOVA yielding a main factor of noise level (F(2)=16.72, p=0.0003) (Figure 4h). Post-hoc comparisons (Tukey HSD) showed that the standard deviation in the low noise condition (mean=2.12, SE=0.62) was significantly lower than in the medium (mean=3.01, SE=0.77; p=0.0062) and high noise (mean=3.53, SE=0.83; p=3.71×10^−5^) conditions.

Finally, we assessed the correlation between slopes of lines fit to *model-generated* responses (i.e., using model-fit means and SDs) and *participants’* responses (Figure 4i). The correlation was strong and highly significant (r=0.97, p=1.48×10^−20^), suggesting that the fit parameters explained the data on a participant-by-participant basis. Overall, the results in Experiment 3 showed that participants’ responses systematically moved closer to the prior with increasing sensory noise (4e-f) and that a Bayesian ideal-observer model largely explained the results (4g-i).

### Anti-Bayesian categorization is driven by image noise but not motor-driven noise

The above results showed that continuous perception across saccades is Bayesian, but categorical perception is anti-Bayesian. What gives rise to this puzzling dichotomy? Since behavior in other categorical tasks often *is* Bayesian^39–43^, our findings are likely more related to the perceptual system we studied than the task structure. In the visual system, object location is signaled via the organization of spatial receptive fields. Receptive fields are continuous from the retina to higher order visual areas^44–47^ and maintain their retinotopic properties even across eye movements^48,49^ and when remapped^50,51,52^. Moreover, neurons in the frontal eye field use continuous tuning to represent object displacements across saccades^53^, the stimulus quantity we probed directly. Thus, the intrinsic organization for processing visual location across saccades is in *continuous* coordinates. If reports of displacement are required in similarly continuous coordinates, the visual system is perhaps well-equipped to use a Bayes optimal strategy. Requiring a *categorical* report of the continuous system might necessitate an alternative strategy. This explanation has two important implications.

First, it implies that anti-Bayesian prior use was driven primarily by the organization of the *visual* system. A potential counterargument is that the Gaussian blob in Experiments 2 and 3 was both the visual object and the saccade target. Blurring it might have added noise to the saccade and consequently the motor prediction (Figure 1a, black arrow), which depends on a copy of the saccade command, in addition to adding noise to the visual input (Figure 1a, red arrow). We did not find evidence of this, however. As a function of Gaussian blur, the standard deviations of saccadic endpoints^54,55^ did not change either parallel or perpendicular to the saccade in either the categorical (Experiment 2; Figure 5a,b) or the continuous (Experiment 3; Figure 5c,d) experiment for humans (Repeated-measures ANOVAs: F(2)=0.32, p=0.7306 in Experiment 2 and F(3)=1.55, p=0.2230 in Experiment 3 for endpoints parallel to the saccade; F(2)=0.14, p=0.8693 in Experiment 2 and F(3)=2.6, p=0.1129 in Experiment 3 for endpoints perpendicular to the saccade). Therefore, uncertainty in the visual input seems to have been the sole factor driving the anti-Bayesian prior use.

**Figure 5.**
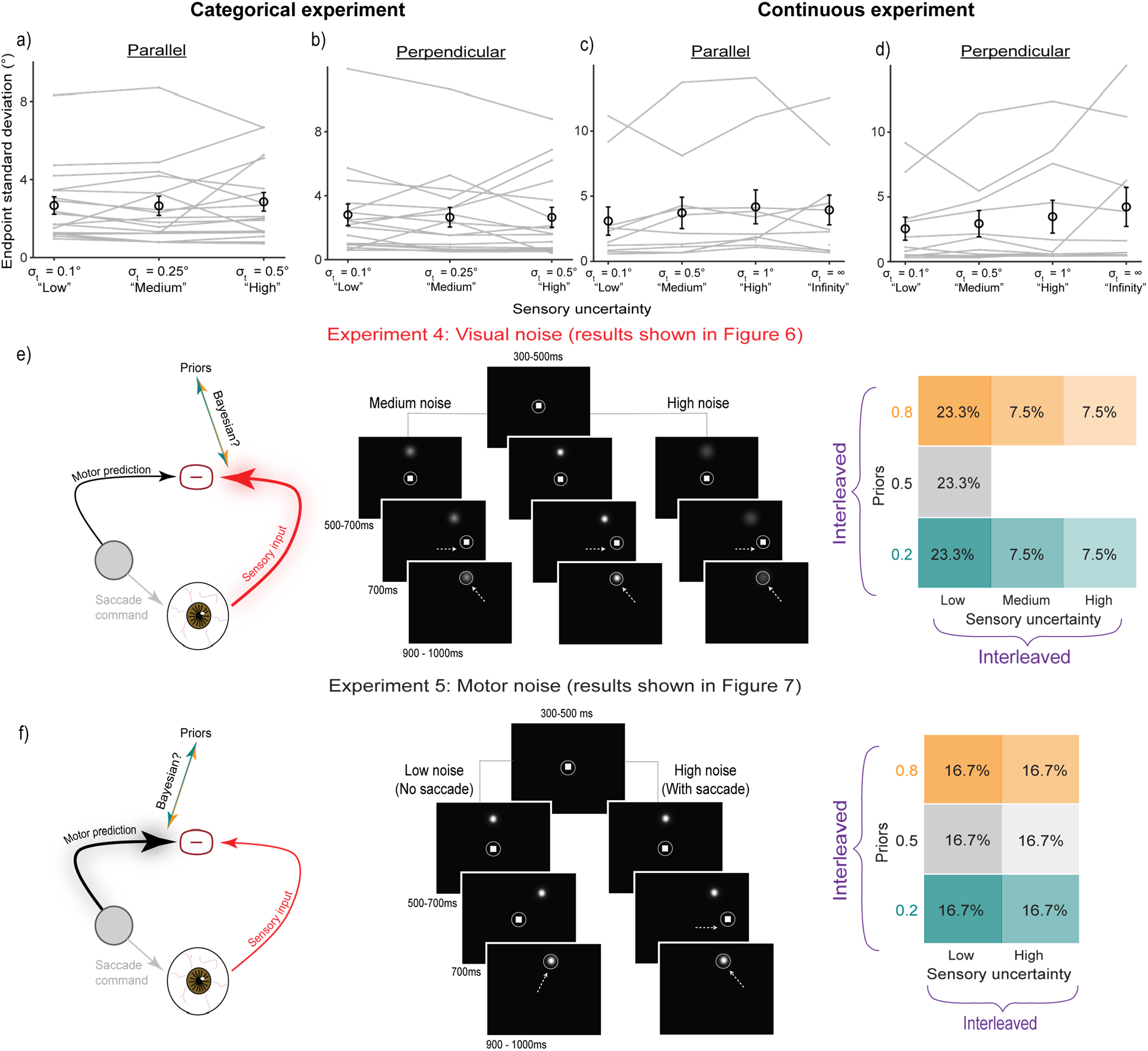
Image noise vs. motor-driven noise. a-d) Saccade endpoint scatter across the three image noise levels in Experiments 2 (Categorical) and 3 (Continuous), as quantified in the directions (a and c) perpendicular and (b and d) parallel to the saccade. (e-f) In monkeys, we separately tested how prior use changes with (e) image noise in Experiment 4 and (f) motor-driven noise in Experiment 5; for each experiment, the rationale (*left*), task events and stimulus configurations (*middle*), and trial breakdown (*right*) are schematized.

Second, the explanation that Bayesian prior use occurs in *continuous* report tasks for the *continuously*-organized visual system implies the converse: Bayesian prior use should occur in *categorical* report tasks for systems having *categorical* properties. Making a saccade is one example. Each saccade poses an inherent, largely categorical sensory uncertainty in the form of saccadic suppression^30,37,56–60^. Visual processing is suppressed when a saccade is made, and not otherwise. This predicts that prior use would be Bayesian to compensate for saccadic suppression.

These considerations suggest a hypothesis that categorical tasks elicit (1) *anti-Bayesian* prior use if the sensory uncertainty is continuous (here, because it is represented in the continuously organized visual system), but (2) *Bayesian* prior use if the sensory uncertainty is categorical (here, because it is due to a saccade being made or not). In Experiments 4 and 5, respectively, we tested these hypotheses. We controlled for motor prediction uncertainty covarying with visual uncertainty by separating the blurred visual stimulus from the saccade target. The experiments used monkeys to permit precise eye position measurements with implanted scleral search coils^61,62^.

*In Experiment 4* (Figure 5e), we selectively manipulated visual uncertainty by varying the width of the Gaussian blob (i.e., the image noise). The structure of Experiment 4 was nearly identical to Experiment 2 in humans (Figure 5e, right): there were three noise levels (low, medium, and high). Low-noise, prior-training trials comprised 70% of all trials, while medium- and high-noise hypothesis-testing trials with neutral jump probability of 0.5 comprised 30% of trials. All trial types were randomly interleaved. *In Experiment 5* (Figure 5f), there were two levels of motor-driven uncertainty. In the “high uncertainty” condition, monkeys made a saccade to a target (to induce saccadic suppression) and reported whether a probe moved or not. In the “low uncertainty” condition, they withheld the saccade (no saccadic suppression) while the probe moved. With- and no-saccade trials at three prior levels each were randomly interleaved (Figure 5f, right).

Results from Experiment 4 replicated the results from Experiment 2: the behavior was anti-Bayesian (Figure 6). As expected from separating the visual probe from the saccade target, there were no significant changes in standard deviations of saccade endpoints across noise levels (Figure S7). Hence prior use was driven by visual uncertainty with no detectable contribution of motor prediction uncertainty. Both animals learned the priors as expected (*P*(*J*)=0.2, 0.5, or 0.8), leading to an upward shift in psychometric functions for the high (*P*(*J*)=0.8) prior and a downward shift for the low (*P*(*J*)=0.2) prior (Figure 6a,b). Quantitatively, intercepts [95% confidence intervals] increased with increasing priors: 0.08 [0.07, 0.10], 0.36 [0.33, 0.39] and 0.57 [0.53, 0.60] for Monkey S (Figure 6c, circles) and 0.18 [0.15, 0.22], 0.33 [0.30, 0.36] and 0.40 [0.31, 0.45] for Monkey T (Figure 6c, triangles).

**Figure 6.**
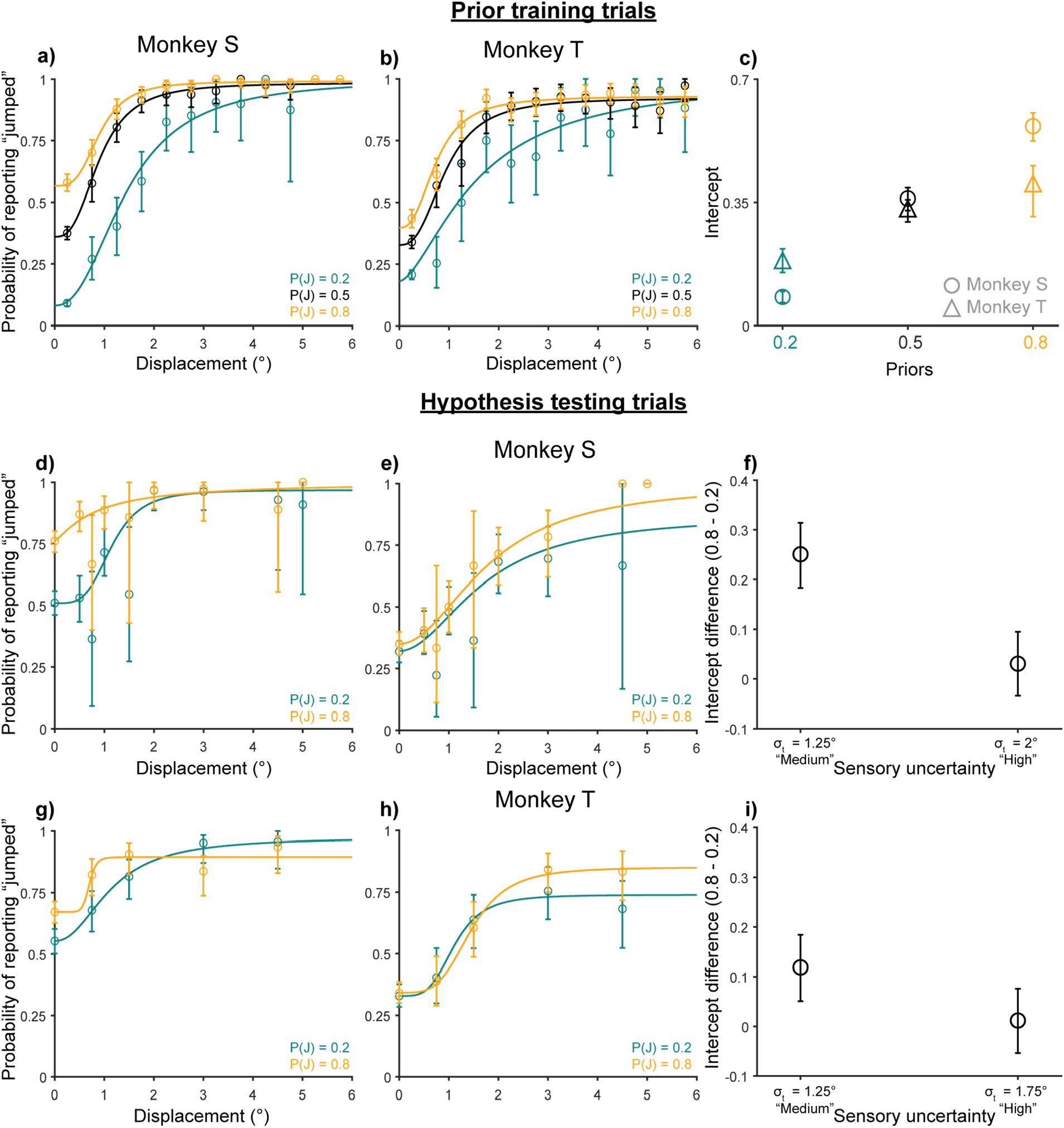
Experiment 4 results: Monkeys were anti-Bayesian for image noise. Monkey S: 10130 trials from 9 sessions. Monkey T: 9958 trials from 18 sessions. 95% confidence intervals bootstrapped from 10000 samples. a-c) Both animals’ performance in prior learning trials in terms of psychometric curves (a: Monkey S; b: Monkey T) and intercept differences (c: both monkeys) matched the predictions of the Bayesian ideal observer model in Figure 2d,e. d-i) Prior use for both monkeys (d-f: Monkey S; g-i: Monkey T) was anti-Bayesian. They showed greater prior use in the medium noise condition (d and g) than in the high noise condition (e and h), as quantified by the intercept differences (f and i).

Also consistent with human participants, prior use *decreased* with increasing noise (Figure 6d-i). Psychometric functions for the 0.2 and 0.8 prior conditions got closer to each other with increasing noise for both Monkey S (Figure 6d,e) and Monkey T (Figure 6g,h), in contrast to the greater separation with noise predicted by a Bayesian model (Figure 3a,b). Intercept differences between the high- and low-prior conditions reflected this collapsing of curves. Monkey S had an intercept difference of 0.25 [0.18, 0.32] in the medium noise condition and 0.03 [-0.03, 0.09] in the high noise condition. For Monkey T, it was 0.12 [0.05, 0.19] for medium noise and 0.01 [-0.05, 0.08] for high noise. Overall, the results showed that both monkeys used their priors less with increasing image noise. Note that the control experiment presented in Figure 3 also selectively varied the width of the Gaussian blob but not the saccade target, replicating the finding that prior use with increasing external, image uncertainty was anti-Bayesian regardless of task structure.

Experiment 5 showed, in contrast, that prior use to account for motor-related noise in the categorical task *was* Bayesian (Figure 7). Since early visual processing and sensitivity to displacements are suppressed around the time of saccades, we simulated motor-driven noise by increasing the standard deviation of the non-jump likelihood distribution in the with-saccade condition relative to the no-saccade condition (σ_NJ_=1° and 0.25°, respectively) while holding the standard deviation of the jump likelihood distribution constant (σ_J_=5°). That is, larger displacements are perceived as “non-jumps” in the with-saccade condition to mimic the saccadic suppression of displacement.

**Figure 7.**
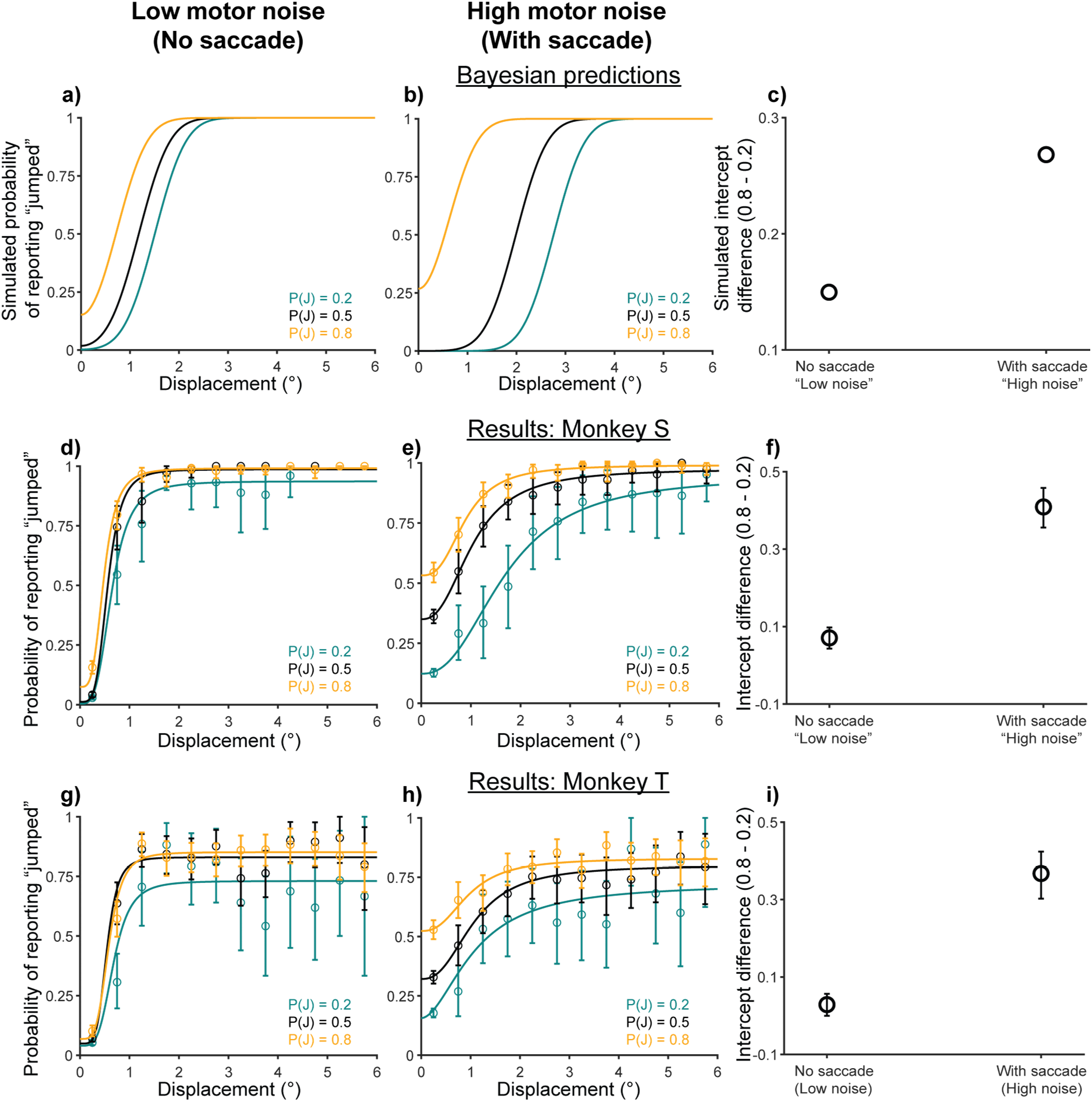
Experiment 5 results: Monkeys were Bayesian for motor-related noise. Overall format similar to Figure 3a-f, with *top row* (a-c) showing the predictions of Bayesian modeling, *middle row* (d-f) illustrating the respective results from Monkey S, and *lower row* (g-i) illustrating the respective results from Monkey T. Unlike for image noise (Figure 6 and Figure 3h-i), the monkeys were decisively Bayesian in their use of priors to compensate for sensory uncertainty introduced by making a saccade.

The Bayesian model predicts that psychometric functions in the 0.2 and 0.8 prior conditions would separate further (Figure 7a,b) and that the difference in intercepts between them would be greater (Figure 7c) with greater motor-driven noise. Results from both monkeys matched these model predictions. Psychometric curves for the different priors showed greater separation when animals made a saccade (Figure 7e: Monkey S; Figure 7h: Monkey T; n=6000 analyzed trials for both animals) than in the condition without a saccade (Figure 7d: Monkey S; Figure 7g: Monkey T). The intercept difference between priors was 0.07 [0.04, 0.10] in the no-saccade condition and 0.41 [0.36, 0.46] when a saccade was made for Monkey S (Figure 7f), and 0.03 [-0.0007, 0.06] and 0.37 [0.30, 0.42] respectively for Monkey T (Figure 7i).

### A discriminative model provides a candidate explanation for anti-Bayesian categorization

In sum, categorical judgments were Bayesian for motor-driven noise but anti-Bayesian for image noise. This leads to the question of how the anti-Bayesian behavior is generated. To address this, we sought to explain two aspects of the results that violated Bayesian predictions. First, prior use decreased with increasing image noise. Second, for human participants, the low prior (teal) curve rose above the baseline (black) curve in Figure 2e, violating the Bayesian prediction of parallel prior psychometric curves (Figure 2d). Although prior curves were parallel for monkeys (Figure 6b,c and Figure 7e,h, respectively), this was after extensive training. Humans performed only single sessions. The early prior-training data from both monkeys were consistent with the human data (Figure 8a,b, teal curves rose above black curves). Therefore, an alternative model would have to explain the disproportionately high “jump” response rate for low priors early in training in addition to the decrease in prior use with increasing visual uncertainty.

**Figure 8.**
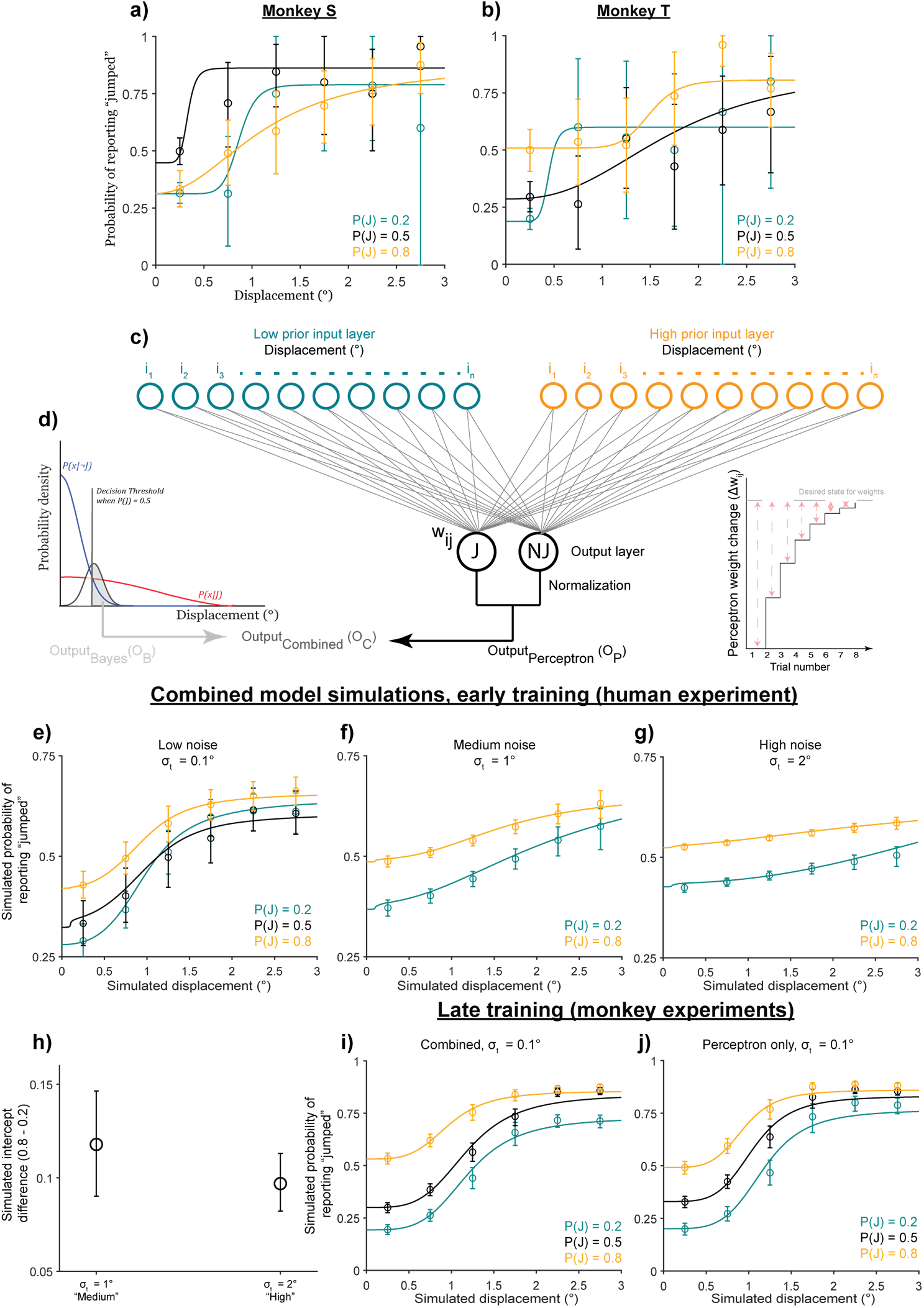
Discriminative learning model. a-b) Early training results (Monkey S trials 1-500 and Monkey T trials 1-350 for each prior). c) Schematic of the discriminative model. Inset schematizes the change in weights with each trial. d) Schematic of the Bayesian model (from Figure 2c). e-f) Psychometric curves simulated under the same experimental conditions as the human experiment for (e) low, (f) medium, and (g) high noise levels. Bins: averaged across 10000 simulations. Error bars: 95% CI. Psychometric curves are averaged across the simulations (blips at small displacements are an artifact of averaging across different inflection points). h) Intercept differences across the medium and high noise conditions. i-j) Late training data for (i) the combined model and (j) the discriminative model alone.

An alternative framework to Bayesian models is discriminative models^4–7^, which directly learn to classify stimuli. For our categorical task, a discriminative model would seek to classify continuous displacements into two categories, “jump” and “no jump.” We set up a simple two-layer neural network to classify displacements (Figure 8c) and simulated its performance under experimental conditions (Figure 8e-j). For details, see *Detailed Methods: Section 4*. To summarize its design, the neural network received continuous displacements in its input layer that were mapped onto “jump” and “no jump” as output categories. The final output was normalized to range from 0-1 to produce a “probability of reporting jumped” given a perceived displacement. The activation of the input layer was determined by the sensory noise on each trial, and there were separate input layers for each prior such that the input-output mapping was prior-dependent. The weights between inputs and outputs were updated in an error-driven manner, rather than a purely associative one^63^, based on work by Gluck and Bower (1988)^64^ who showed that error-driven learning leads to a disproportionate overweighting of infrequent events. This is equivalent to the Perceptron learning rule^65,66^. Since errors are large early in training, weight changes are relatively large, too (left side of Figure 8c, inset; red dashed arrows depict the error). Over time, as the errors decrease, the weights asymptote towards the desired state (right side of Figure 8c, inset). For infrequent events such as large displacements in the low prior condition early in training, weight changes are relatively large. Taking a snapshot of performance at this stage results in an apparent overweighting of their contribution to the response rate (Figures 2e and 8a,b). Once weights approach the desired state late in training, events contribute proportional to their relative frequencies, and the psychometric curves become more parallel to one another (Figures 6b,c and 7e,h).

We simulated model performance under experimental conditions. First, we looked at early prior learning. We simulated early learning using the same task structure and closely-matched parameters as Experiment 2. As expected, the outputs recapitulated the disproportionately high response rate for large displacements in the low prior condition (Figure S4b, teal curve). However, it also downweighted the infrequent *small* displacements in the high prior condition (Figure S8b, orange curve). To account for this, we considered that reports in the categorical task result from a *combination* of a discriminative (Figure 8c) and a Bayesian (Figure 8d) model. The Bayesian prior use for high saccade-driven uncertainty raises high prior intercepts (Figure 7e,h) and thus could compensate for the downweighting. Therefore, we next simulated a linearly weighted combination of outputs from the discriminative and Bayesian models in the prior training condition (Figure 8c,d, bottom; see *Detailed Methods* for details). Combining the outputs of the Bayesian and discriminative models at relative weights of 0.1 and 0.9 respectively generated the data in Figure 8e, where the pattern of the curves matched data from human participants well. Next, adding medium and high noise to the visual input of the discriminative model (but holding visual noise constant in the Bayesian model) caused psychometric curves to move closer to each other with increasing noise (Figure 8f,g), as quantified by the downward trend in high-low prior intercept differences (Figure 8h). Confirmation that visual uncertainty alone flattened psychometric curves in Figure S8b. Finally, we tested the prediction that prior curves become parallel once weights approach a relatively stable desired state for all input units (Figure 8c, inset, right) by letting the model run for 5000 trials. Data from trials 3000-5000 for the combined (discriminative + Bayesian) model (Figure 8i) and for the discriminative model alone (Figure 8j) support this prediction.

In summary, the combined model recapitulated both the surprising trade-off between priors and noise and the long-term learning effects that were unexplained by a Bayesian ideal observer model alone. This demonstrated that a discriminative learning rule provides a feasible explanation for the anti-Bayesian categorization of object displacement across saccades.

## Discussion

We found that human participants were Bayesian for continuous reports of object displacement across saccades but anti-Bayesian for categorical reports. Further investigation in monkeys showed that the anti-Bayesian effect was primarily due to external, image noise rather than motor-driven noise. The use of a Perceptron-like, discriminative learning rule provides a candidate explanation for anti-Bayesian performance in the categorization task.

Limitations of the experimental design and modeling choices should be considered while interpreting the results. First, it is possible that the anti-Bayesian result is a consequence of how we conceptualized parameters (e.g., the prior, or sensory noise) in the categorical Bayesian ideal observer model. For example, for continuous tasks, it has been shown that if the sensory likelihood is asymmetric in a way that can result from assumptions of efficient sensory encoding, then the outcome of a Bayesian decoding process can be seemingly anti-Bayesian^67^. Of course, alternative parameters might predict the surprising results. However, we chose simulation parameters to closely map onto experimental parameters and mimic empirical phenomena such as saccadic suppression. As a result, the model captures both the Bayesian trade-off with motor-driven noise and the anti-Bayesian trade-off with visual noise. To our best estimate, there was no simple set of alternative parameters that did so parsimoniously.

Second, we limited the simulation of motor-induced noise in Experiment 5 to just one phenomenon, i.e., saccadic suppression. We did this by increasing the width of the non-jump likelihood, 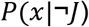. We focused on saccadic suppression since it is largely a categorical form of uncertainty that is present when a saccade is made, and not otherwise. It was thus sufficient for testing our hypothesis. However, there are other ways in which saccades influence vision both at the level of behavior and neurons. Such effects include compression of space toward the saccade target^68–71^ or the shifting or smearing of visual receptive fields around the time of saccades^51,52^. Indeed, the magnitude of saccadic suppression may vary with saccade amplitude^72^ and the direction in which the probe moves relative to the saccade vector^31,53^. Our results do not preclude the inclusion of additional, fine-grained motor-induced phenomena into the normative model, and the resulting predictions would be testable. Another consideration for Experiment 5 is that in the no-saccade condition of the task, animals fixated a central square for the duration of a trial. We did not prevent the animal from making fixational eye movements such as microsaccades^73^, and saccadic suppression may occur around the time of microsaccades^74–77^. Although we did not control for microsaccades, the stimulus displacement was not timed to microsaccade onset in no-saccade trials as it was to saccade onset in with-saccade trials. On average, therefore, the influence of (micro)saccadic suppression should be quite low in the no-saccade condition.

Finally, we limited the scope of the discriminative and combined models to provide a candidate alternative to the *categorical* Bayesian model with minimal additional assumptions. This leaves some unexplained patterns in the data, e.g., overall intercepts across priors decrease with increasing noise for humans (Figure. 3d,e) and monkeys (Figure 6d,e,g,h) but not for the model (Figure 8e-g). Similarly, for our experimental-like parameters, the model does not capture the complete collapse of prior curves with increasing noise. The model may be extended, however, to include additional components such as Bayesian integration in the *continuous* input layer (from Experiment 3) to better explain the data. Assumptions about how the components combine may additionally be testable too.

The overall pattern of results in our study poses a fundamental question: what determines the use of Bayesian vs. discriminative models for perception? Despite recent efforts to acknowledge the contribution of both Bayesian and non-Bayesian^78,79^ models to perception^11–13^, the field lacks a synthesized, theoretical account of when Bayesian models are used and when they are not. As we noted while motivating Experiments 4 and 5, our results suggest a link between the inherent neural organization of sensorimotor systems and Bayesian behavior. Further clarification of this link would allow our understanding of each to constrain and advance our understanding of the other.

Finally, we expect that the conclusions of this study extend beyond the oculomotor system. Accounting for self-movement is an issue for almost all sensory modalities, and the integration of movement and sensory signals for active perception has been observed widely in the brain^80–83^. Understanding the relative contributions of Bayesian and discriminative computations to active vision may guide future studies on how expectations, self-movement, and external sensory information combine for more general forms of active perception.

## Acknowledgments

We would like to thank Jessi Cruger for caring for the non-human primates used in this study and Seth Madlon-Kay for feedback and discussions. The work was supported by the Duke Institute for Brain Sciences Research Incubator Award (2020).

## Author Contributions

Conceptualization, D.S., J.M.P. and M.A.S.; Methodology, D.S., J.M.P., and M.A.S.; Software, D.S. and J.M.P.; Formal Analysis, D.S. and J.M.P.; Investigation, D.S.; Resources, M.A.S.; Data Curation, D.S.; Writing – Original Draft, D.S. and M.A.S.; Writing – Review & Editing, D.S., J.M.P., and M.A.S.; Visualization, D.S. and M.A.S.; Supervision, M.A.S.; Project Administration, D.S. and M.A.S.; Funding Acquisition, D.S., J.M.P, and M.A.S.

## Detailed Methods

### Methods Overview

The following Methods section is split into 4 sub-sections. Section 1 includes the methods for the psychophysics experiments run on humans. This includes an initial experiment to select the stimulus to be used as a sensory noise manipulation (Experiment 1), an experiment testing the trade-off between categorical priors and sensory uncertainty (Experiment 2), and an experiment testing the trade-off between continuous priors and sensory uncertainty (Experiment 3). Section 2 details the methods for the experiments run on rhesus macaques. This section includes the experiment to isolate the trade-off between a categorical prior and visual uncertainty alone (Experiment 4), between a categorical prior and motor uncertainty alone (Experiment 5), and a control experiment. Section 3 includes a description of the data preparation and analysis measures used throughout the manuscript. Section 4 includes a detailed description of the computational models used in the manuscript.

#### Section 1. Human Psychophysics

##### Materials and Paradigm

45 adult volunteers with normal or corrected-to-normal vision participated in the experiments. All procedures were explained verbally to participants beforehand and written, informed consent was obtained. Participants were paid $12/hr. and informed that participation was completely voluntary. All procedures were performed in accordance with protocols approved by the Duke University Institutional Review Board.

Participants sat alone in a darkened room in front of a monitor with their head stabilized using a chin- and forehead-rest. The monitor was positioned at 60cm from the center of the head. Experiments 1 and 2 were displayed on a CRT monitor (Accusync 120) at 120Hz. Experiment 3 was displayed on a Dell LCD monitor at 60Hz. The experiment was written in and displayed using Presentation (Neurobehavioral systems). Monocular eye position was recorded with an eye-tracking software developed by Matsuda et al. (2017)^84^.

Participants performed a modified Saccadic Suppression of Displacement task^37^. On each trial, a fixation cross first appeared near the center of the screen. Once participants had acquired and maintained fixation for 500ms, a saccade target appeared at one of two average positions relative to the center of the screen: 10° or −10°. A target at 10° appeared in the right half of the screen while a target at −10 appeared in the left half of the screen. Additionally, on every trial, the position of the target and fixation cross were both jittered by −0.5 to 0.5 degrees relative to the average position to mitigate the confounding effects of adaptation to either a constant saccade amplitude or a constant distance between the target and the edge of the screen. The fixation cross then disappeared for 500ms, and an auditory cue was presented to signal to participants that they were allowed to make a saccade to the target. If fixation was broken before the auditory cue, the trial was aborted and a new one began immediately. Saccade initiation (defined as the time the eye left a window of 2 deg. relative to the fixation cross) triggered target displacement. In Experiments 1 and 2, participants provided a binary, categorical report on whether they had perceived the target as having moved or not. The target remained on the screen for 500ms after it was displaced, after which it was replaced by a response prompt screen (5 = moved, 0 = remained stationary). In these experiments, target displacement was drawn from overlapping Gaussian distributions designated as the “movement” and “non-movement” distributions (Figure S1). Displacements were drawn from overlapping, rather than distinct, distributions to ensure that the solution to the task was probabilistic. On trials where the target moved, the displacement was drawn from a relatively broad Gaussian distribution centered around 0 (μ=0°, σ=1.5°). On “no movement” trials, the displacement was drawn from a very narrow Gaussian distribution centered around 0 (μ=0°, σ=0.017°). A positive displacement meant that the target moved rightward, and a negative displacement meant it moved leftward.

In Experiment 3, they provided a continuous report of the target’s postsaccadic location. For this study, the target stayed visible for 100ms after displacement and was then replaced by a screen where the mouse cursor (shaped “+”) was placed at the center of the screen and restricted to the horizontal meridian. Participants could then move the mouse cursor to where they perceived the target as having landed.

##### Preliminary Experiment for visual uncertainty manipulation

The goal of Experiment 2 was to test the use of prior expectations with increasing sensory noise. The main experimental variables were thus priors and sensory noise. We planned to cue priors (jump probability) by the color of the fixation cross and train participants on this association using performance-based feedback. We were confident that this prior manipulation would be successful since it was only a slight variation from previous work in the lab^36^. As a first step, therefore, we sought to confirm that our sensory noise manipulation reliably induced uncertainty. 9 human participants completed at least 100 trials each in 8 experimental conditions: 4 candidate stimuli at two uncertainty levels each. The probability of target movement across all stimulus conditions was 0.5.

The 4 possible stimuli were:

1. A Gaussian cloud of 20 white squares (0.25° x 0.25°) where the uncertainty corresponded to the standard deviation of the cloud (low uncertainty = 0.0625° and high uncertainty = 0.25°).
2. Arrows (1° long with 0.5° width) that either pointed in the direction of the jump (congruent) or in the opposite direction (incongruent). We predicted that incongruent jumps might induce greater uncertainty and decrease discriminability.
3. Squares (0.5° x 0.5°) at two levels of contrast (low uncertainty = 0.784 and high uncertainty = 0.294).
4. Gaussian “blobs” (isoluminant, Gaussian distributions of light) uncertainty corresponded to the standard deviation of the blob (low uncertainty = 0.19° and high uncertainty = 0.47°).

The results of this experiment would determine how we manipulated the sensory uncertainty of the stimulus in all other experiments.

##### Experiment 2: Trade-off between binary prior and sensory uncertainty in humans

We trained participants on the prior, cued by the color of the fixation cross, using performance-based feedback. They were told whether their responses were correct or incorrect on each trial using an image of a smiling or frowning face, respectively. Based on the results from Experiment 1, we chose the isoluminant Gaussian blob as sensory uncertainty manipulation. The target was grayscale on every trial; only its width given by the standard deviation changed. The target had one of three possible standard deviations for the whole experiment – 0.1 deg. (“low noise”), 0.25 deg. (“medium noise) or 0.5 deg. (“high noise”).

Twenty participants completed a total of 1300 trials each. Trials were presented in 100-trial blocks. For all participants, the first block was a baseline block where the color of the fixation cross was white, and the target moved on 50% of the trials. In the next 6 blocks, the fixation cross was either green or red, and vice versa for last 6. Each of these fixation colors was associated with one of two probabilities of target movement (0.9 or 0.1). The order of the two prior conditions and color-probability associations were counterbalanced across participants. As in Experiment 1, displacements were drawn from a relatively broad Gaussian distribution (μ=0°, σ=1.5°) on “movement” trials and from a narrow Gaussian distribution (μ=0°, σ=0.017°) on “non-movement” trials to ensure that the solution to the task was probabilistic. Thus, the optimal solution to the task was to learn the probability that any given displacement was drawn from the “movement” distribution relative to the “non-movement” distribution. In conditions with a biased prior (0.9 or 0.1), the optimal solution would be to weight this relative probability by the appropriate prior. In other words, the optimal solution to this task is the Bayesian solution.

For 70% of the trials in blocks 2-13, the target had the lowest noise (standard deviation of 0.1°) and the probability of target displacement conformed to the experimental prior, i.e. 0.9 or 0.1. These 70% of the trials were considered “training trials” where the intended prior was reinforced and maintained. The other 30% of trials were “testing trials”, where we tested the hypothesis that participants would use their learned prior more when the evidence was relatively uncertain. On these trials, the target had either medium or high sensory noise. Additionally, both to isolate the effects of a learned, color-associated expectation on performance and to mitigate the possibility that our sensory manipulation affected participants’ representation of the prior, the testing trials comprised a neutral condition where the target had a 0.5 probability of moving, but the fixation color cuing the prior was the same as the rest of the block. Training and testing trials were randomly interleaved. To preserve a sense of experiential continuity across the experiment, 5% of the targets in block 1 had a standard deviation of 0.25 deg. (“medium-noise”) and the remaining 5% had a standard deviation of 0.5 deg. (“high-noise”). Data from these trials are not analyzed.

##### Experiment 3: Trade-off between continuous prior and sensory uncertainty in human participants

Fourteen human volunteers participated in Experiment 3. We tested the hypothesis that the visual system uses Bayesian Inference to determine the *continuous* displacement value of objects across saccades. The overall paradigm was similar to Experiments 1 and 2. The critical difference was that the target was displayed post-saccadically for a limited period (100ms) and participants provided a continuous report of where they had perceived it as having landed. Participants fixated a central cross and upon being cued, made a saccade to a target located at either 10° or −10°. The target was displaced horizontally during the saccade, displayed in its new location for 50ms, and then replaced by a response screen. The response screen consisted of a mouse cursor that started out in the center and was restricted to the horizontal meridian to ensure that participants were solving a one-dimensional problem. Participants were required to drag the mouse cursor to the location where the target had landed and click to submit their response.

Participants completed a total of 1000 trials each. We first trained participants on the prior for 600 trials and then tested the use of this prior with increasing sensory noise. As in Experiments 1 and 2, the target was a Gaussian blob and we manipulated sensory uncertainty by varying its width. The prior was a *continuous* Gaussian distribution of displacements, rather than a categorical prior indicating the probability of object displacement. Throughout the experiment, displacements were drawn from a Gaussian distribution with mean 0° and standard deviation 1°. Participants were trained on this prior in the first 600 trials with performance-based feedback. After they submitted their response, the target appeared in its correct postsaccadic location for 500ms. To indicate their degree of correctness, the color of this feedback target ranged continuously from green (correct) to red (incorrect by more than 2°). Targets in this training phase trials had a standard deviation of 0.1° (“low noise”).

In the remaining 400 trials, participants underwent a “testing” trials phase they were provided no feedback. These trials had one of 3 noise levels - 0.1° (“low noise”), 0.5° (“medium noise”), and 1° (“high noise”). Further, throughout the experiment, in 20% of the trials, the target did not appear post-saccadically. We call these “infinite noise” trials. All 4 noise levels were randomly interleaved throughout the testing phase. All data shown in Figure 3 comes from the testing phase of the experiment. We used participants’ performance in the infinite noise condition as a means to evaluate how well they learned the prior in the training phase.

#### Section 2. Rhesus Macaque Psychophysics

##### Materials and Paradigm

Two Rhesus macaques (Monkey S and Monkey T, both males) were run in a modified Saccadic Suppression of Displacement paradigm, similar to the human participants. Animals were brought into the lab in custom-made chairs (Crist Instruments, Hagerstown, MD) and their heads were stabilized using a head-post that attached to both the chair and a surgically implanted socket (Crist Instruments, Hagerstown, MD) on the skull. The socket was implanted in an aseptic surgical procedure with the help of ceramic screws and acrylic. Eye position was measured using a surgically implanted scleral search coil^61,62^ in one eye. All surgical and experimental procedures were performed in accordance with protocols approved by the Duke Institutional Animal Care and Use Committee.

In a typical experimental session, the animals performed the behavioral task in a dark experimental rig. They were positioned 60cm from an LCD monitor (1920 × 1080, 144Hz). To dissociate external sources of sensory noise from internal, motor-driven sources, the saccade target was dissociated from a visual probe (a Gaussian blob) which was displaced intrasaccadically on some trials. In the human experiments, the Gaussian, visual probe (same as the saccade target) always appeared in one of two locations on the screen and only moved horizontally. For Experiments 4 and 5 in monkeys, it could appear in one of 4 locations, ±10° horizontally or ±10° vertically. The saccade target was always positioned along the orthogonal cardinal direction (e.g. if the probe appeared at ±10° horizontally, the saccade target would be at ±10° vertically), and the probe was displaced in a direction parallel to the saccade vector. For the control experiment (with results presented in Figure 3), we simultaneously recorded from neurons while the animals performed the sessions (neural data not presented in this manuscript). Since we placed the probe within the mapped receptive field of the neuron, the probe appeared in a different location during each session.

On each trial, a fixation square (1°x1°) first appeared at the center of the screen. After fixation had been acquired and maintained for a randomized duration of 300-500ms, the visual probe appeared at one of the 4 locations on the screen for 500-700ms. The monkey was required to maintain fixation on the central fixation square for that duration, after which the fixation square was replaced by the saccade target (1°x1°) indicating to the animal they could make a saccade. Saccade initiation (defined as the time the eye crossed a threshold set at 20% of the saccade length, i.e., 2°, in the direction of the saccade) triggered the displacement of the probe on some trials. The probe was displaced in a direction parallel to the saccade. Animals were further required to maintain postsaccadic fixation for 700ms after which the saccade target was replaced by a white cross in the same location. To report that the probe had moved during the saccade, the monkey was required to make a saccade to the probe within 500ms and then fixate on it for 400ms. To report that it had remained stationary during the saccade, the monkey had to remain fixated on the cross for 1000ms. The precise timing of stimulus presentation was verified with a photodiode taped to the top left corner of the monitor, where a white square (invisible to the monkey) was flashed within the same frame as the measured stimulus.

Displacements were drawn from relatively broad and narrow Gaussian distributions in the “movement” and “non-movement” conditions, respectively. Both distributions had a mean of 0°. The “movement” distribution had a standard deviation of 2.5° and the “non-movement” distribution had a standard deviation of 0.2°. Positive displacements were either rightward or upward, and negative displacements were leftward or downward. Priors were cued by the color of the fixation and target squares. For monkey S, green squares meant that the probe had a 0.2 probability of being displaced while magenta squares indicated a 0.8 probability of displacement. For monkey T, blue squares were associated with a 0.2 probability of displacement while orange squares were associated with a 0.8 probability of displacement. Animals were trained on priors over multiple sessions using performance-based feedback like human participants.

##### Experiment 4: Trade-off between categorical priors and *visually-driven* sensory uncertainty

To measure performance as a function of external sensory uncertainty, the visual probe in Experiment 4 was a Gaussian “blob” with one of three possible standard deviations: 0.5° (“low noise”), 1.25° (“medium noise”), and 2° (“high noise”) for Monkey S and 0.5° (“low noise”), 1.25°(“medium noise”), and 1.75° (“high noise”), for Monkey T. The relative frequencies of all trial 7 types (2 priors x 3 noise levels + baseline) were the same as in the categorical experiment for humans (Experiment 2). Baseline trials with white squares and 0.5 probability of displacement all had “no noise” visual probes. In the 0.2 and 0.8 prior conditions, 70% of trials had no noise and conformed to the displacement probability indicated by the prior. The remaining 30% of trials with low and high noise comprised a neutral “test” condition with a veridical jump probability of 0.5. All 7 trial types were randomly interleaved.

##### Control experiment with valid prior statistics for all noise levels

We also analyzed data from a control experiment where the jump probability was the same for all noise levels. In this experiment too, we selectively manipulated external visual noise. Visual noise levels were the same as those in Experiment 4.

##### Experiment 5: Trade-off between categorical priors and *motor-driven* sensory uncertainty

To measure performance as a function of internal, motor-driven sensory uncertainty, we added a condition to the experiment where monkeys did not make a saccade. They remained fixated in the center while the Gaussian, visual probe was displaced. This no-saccade condition served as the “low motor noise” condition and was compared to a “high motor noise” condition where animals made a saccade. The temporal structure of the no-saccade trials was identical to the trials with a saccade. No-saccade trials were implemented by assigning the location of the “saccade target” to be the same as the fixation square. There were three prior conditions: 0.2, 0.5, and 0.8. Colors indicating the priors were the same as in Experiment 4. The visual probe had a standard deviation of 0.5°, the lowest noise condition, for all trials. All 6 trial types (3 priors x 2 noise levels) were randomly interleaved.

#### Section 3. Data preparation and Analysis Measures

##### Data preparation

Data from individual trials were analyzed *post hoc* to confirm that the visual probe landed in its displaced location before the end of the saccade. The saccade end time was defined as the time at which the eye velocity dropped below 0.04°/ms. For human participants, the time at which the target jump command was sent was recorded for each trial. Trials with a recorded jump time greater than 1 whole frame (8.33ms for Experiments 1 and 2, and 16.7ms for Experiment 3) before the detected end of the saccade were excluded from analysis. Participants for whom at least 90% of all trials did not meet this criterion were excluded from analyses entirely. No participants were excluded in Experiment 1, three participants were excluded from Experiment 2, and three participants were excluded from Experiment 3. For the macaque experiments, we used a photodiode to verify the exact timing of stimulus presentation. Note that the timestamp from the photodiode indicated the presentation of a white square at the top left of the screen and the monitor refreshes frames as a raster. We verified the maximum duration of a frame as being 7ms from top left to bottom right using a second photodiode. Since the probe was presented at various locations on the screen, we set the most conservative criterion such that the photodiode timestamp had to be at least 7ms before the detected end of the saccade. Individual trials that did not meet this criterion in the macaque data were excluded as well.

##### Psychometric curves and prior use

All data were analyzed using MATLAB (Mathworks, Inc.). For Experiments 1, 2, 4, and 5, Psychometric curves were fit to binary responses using a 4-parameter logistic regression model:

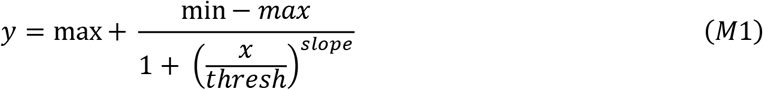

where *x* is the absolute value of the presented displacement, *y* is the value of the psychometric function, *min* is the minimum value of the function (i.e., *y* at *x*=0), *max* is the maximum value, *thresh* is the inflection point, and *slope* is the slope of the psychometric function. *min, max, thresh*, and *slope* terms were fit to binary data by minimizing mean squared error.

For all analyses in the main manuscript, we used the intercept of the psychometric curve as a measure of prior use in these experiments for statistical tests, and for comparison with the predictions of the categorical Bayesian model. For human participants and in the low-noise, prior training trials for macaques, displacements were drawn from continuous distributions. In these conditions, we used the value of *min* as the intercept. For the medium and high noise hypothesis testing trials in macaques, displacements were discretized. There was a displacement = 0 condition. In these conditions, the intercept is simply the proportion of “moved” responses in the displacement = 0 condition.

We repeated all the analyses of prior use presented in the manuscript using a measure from Signal Detection Theory^38^, the Criterion. Criterion provides an alternative measure of bias in responses (i.e., a translational shift in psychometric curves). It is given by:

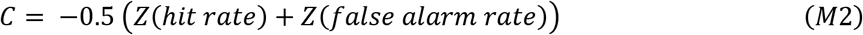

where the hit rate is the proportion of “jumped” responses on trials in which the probe truly moved, and the false alarm rate is the proportion of “jumped” responses on trials in which the probe did not move. The results using Criterion replicate the findings in the main manuscript.

For all statistical comparisons, the assumption of normality was first tested for each sample using a Kolmogorov-Smirnov (KS) test. If met, then we used a parametric comparison such as an ANOVA or a t-test. Otherwise, the equivalent non-parametric test was used.

#### Section 4. Model Simulations and Fitting

##### Categorical Bayesian Ideal Observer Model

The ideal observer makes a probabilistic decision about binary variable, *J*, indicating whether the target jumped or not. ¬*J* indicates that the target did not jump. Since the true displacement is experimentally drawn but not available to the observer, they make this decision given the *perceived* displacement, 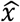. The decision is based on the relative probabilities of the target having jumped or not jumped given the perceived displacement:

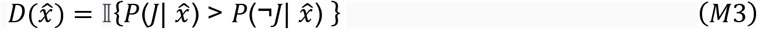

where 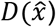 is the decision given the perceived displacement, 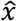, and is determined by a binary indicator function, 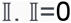 (no jump) if the condition in braces is not met. Otherwise, 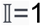 (jumped); 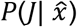 is the probability that the probe jumped given 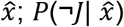 is the probability that the probe did not jump given 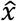.

Using Bayes’ rule for the condition within braces in Eq. M3,

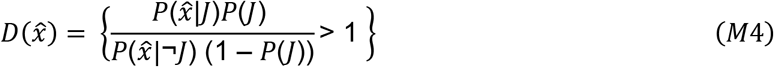

The simulated decision of the ideal observer, however, must be compared to the responses of participants. We do not have access to participants’ perceived displacement, but instead can only infer their decision given the true experimental displacement, *x*. We assume that the perceived displacement is a Gaussian random variable where the mean is the true displacement, and its variance given by the width of the blob on that trial:

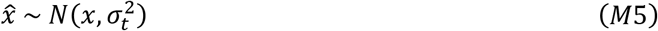

where *σ_t_* is the variance of the target.

The decision given the true displacement can thus be modeled as:

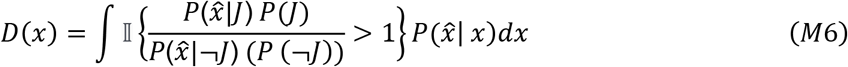

That is, the decision given the true displacement, *D*(*x*), is the integral of the perceived displacement distribution that falls above the point at which the indicator function, 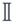, is non-zero.

Based on the distributions used in the experiment (Figure S1),

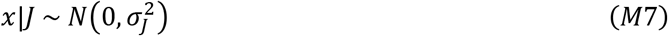

and

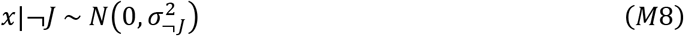

Since 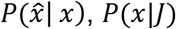, and *P*(*x*|¬*J*) are all Gaussian distributions, we can integrate over *x* such that:

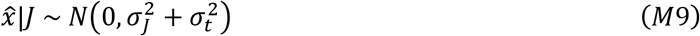

and

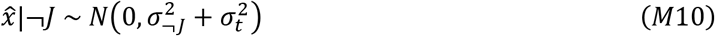

Thus, the expression inside the indicator function in Eq. M6, when replaced with the appropriate Gaussian probability density functions, equals:

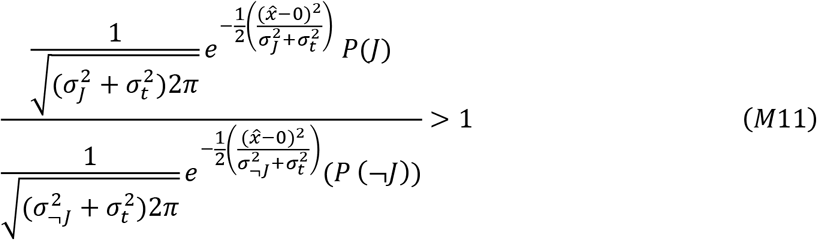

Taking the log on both sides provides the condition under which the indicator function is greater than 0,

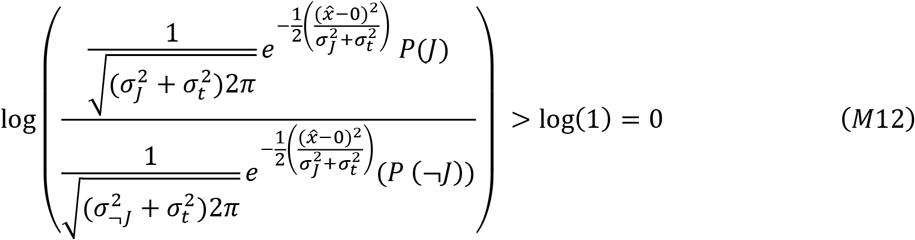

That is,

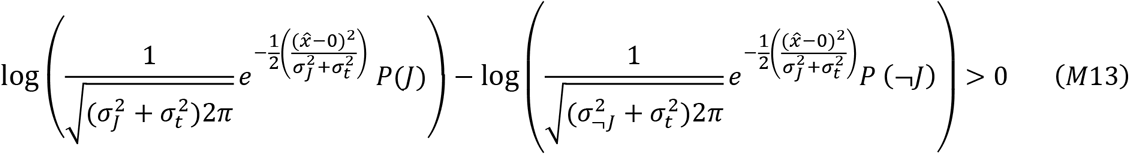

Rearranging terms to solve for 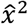, we find that the indicator function is > 0 when 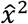 is greater than a criterion value, 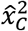, defined as:

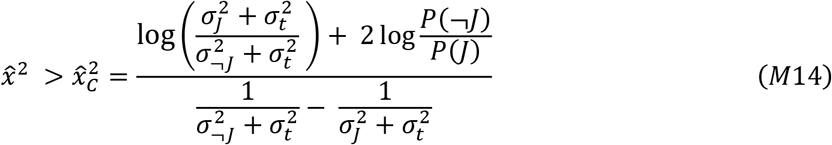

Since 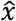 is a Gaussian random variable, 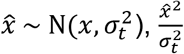 is a non-central chi-square random variable, 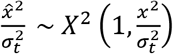. Thus, the decision, *D*(*x*), can be modeled as the integral of a non-central chi-square distribution that lies above the criterion, 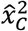. That is,

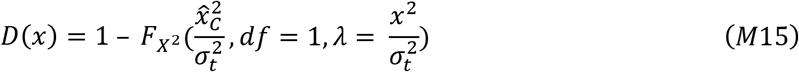

where *F*_*X*^2^_ is the cumulative distribution function of *X*^2^ with degrees of freedom, *df* = 1 and gamma, 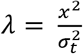 up to 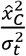.

##### Continuous Bayesian Model: Simulation and Fitting

We first simulated the responses of a Bayesian ideal observer in Experiment 3. The ideal observer infers the perceived displacement as a reliability-weighted combination of the sensory likelihood and prior distributions:

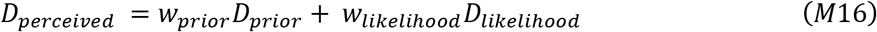

where *D_perceived_* is the mean of the inferred posterior distribution, *w_prior_* is the weight assigned to the prior, *D_prior_* is the mean of the prior distribution, *w_likelihood_* is the weight assigned to the likelihood, and *D_likelihood_* the likelihood distribution. When both *D_prior_* and *D_likelihood_* are Gaussian distributions, the weight terms are given by:

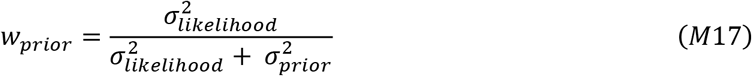

and

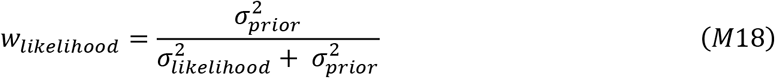

That is, the more reliable (i.e., less variable) estimate is weighted higher. This reliability-weighted inference is additionally the Bayes optimal estimate because the variance of the estimate, 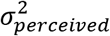, is lower than the variance of both the prior and the likelihood distributions:

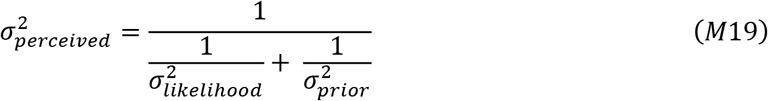

We simulated the final response as the mean of the posterior distribution, i.e., its maximum value. The values of the parameters used to simulate ideal observer responses were the same as the ones used in the experiment.

We then minimized squared error between each participant’s responses and those of the Bayesian ideal observer model to identify the best-fit values for their internal prior and likelihood distributions. Best-fit parameters were identified on a participant-by-participant basis. Parameter optimization was performed using Matlab’s *fmincon* function.

##### Discriminative Model

The goal of the discriminative model was to capture two aspects of the results in the categorical tasks in humans and monkeys that were otherwise unexplained by the Bayesian ideal observer model. First, psychometric curves across prior conditions collapsed together with increasing sensory noise. Second, low prior psychometric curves had a disproportionately high slope early in training but not late in training. We found that designing a simple, two-layer neural network model with an error-based (Perceptron) learning rule and simulating its performance under experimental conditions recapitulated the unexplained patterns.

##### Model structure

The input layer consisted of units representing continuous displacements and the output layer had two units: “jump (J)” and “no jump (NJ).” For ease of computation, continuous input displacements were discretized into bins of 0.1°. Displacements ranged from 0-7.5°. That is, there were 75 input units for each network. Sensory noise was simulated as a Gaussian distribution of input unit activation, truncated at the two ends of the input range (0 and 7.5), such that the total activation of input units was always 1. On each trial, the distribution was centered on the true displacement for the trial and the width of activation was determined by the sensory noise level. Each input unit was connected to both output units. The activation of each output unit was the *weighted* sum of inputs, i.e.:

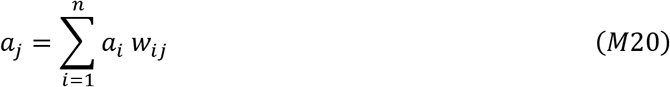

where *a_j_* is the activation of the output unit, j; *a_i_* is the activation of the output unit, i; and *w_ij_* is the weight of the connection between input unit, i, and output unit, j. The final output on each trial was the normalized activation of the “jump” and “no jump” output units such that the output for each unit was bounded between 0 and 1:

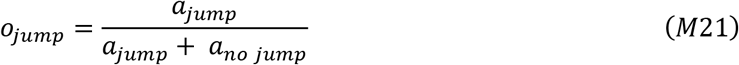

and

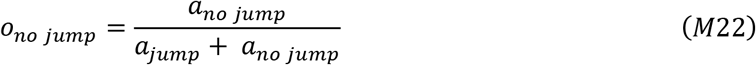

where *o_jump_* is the final output of the “jump” unit, *o_no jump_* is the final output of “no jump” unit, and *a_jump_* and *a_no jump_* are activations of the “jump” and “no jump” output units, respectively.

The “knowledge” of the two categories is stored in the weights between the input and output units, and the learned, category boundary takes the form of a psychometric function reporting the probability that the target “jumped” given an input displacement. In other words, the shape of the psychometric function is determined by the activation of the inputs and the weights between inputs and outputs. Since psychometric curves have different shapes across priors at the same point in training (e.g., Figure 2e), we assumed that the connections between inputs and outputs (and their corresponding weights) are *prior-dependent*. This is equivalent to the idea that distributions are learned separately across cue color contexts. We simulated the prior-dependence of the input-output relationship by simply setting up separate sets of inputs for each prior (Figure 8c).

We next considered how the weights between inputs and outputs might be updated. One possibility was that they are updated by a simple Hebbian-like associative learning rule^59^ where weights between two units are updated in a manner proportional to their activation. We chose a slight variation of this rule based on work by Gluck and Bower^60^, who showed that an *error*based learning rule, rather than a purely associative learning rule, leads to a disproportionate overweighting of infrequent events early in training.

The learning rule is given by:

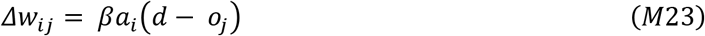

where Δ*w_ij_* is the change in weights between input unit, i, and output unit, j; *β* is the learning rate, *a_i_*, is the activation of the input unit, i; *o_j_* is the final output of unit j, and d is the desired state of output unit, j. The term (*d* – *o_j_*) is therefore the error between the current output of the model and the desired state determined by feedback on each trial. In summary, the change in weights or learning on each trial is proportional to the activity of the input and the error of the model on that trial. This is equivalent to the Perceptron learning rule^61,62^. Since errors are large early in training, weight changes are relatively large too (Figure 8c, inset; red dashed arrows depict the error). Over time, as the errors get smaller, the weights asymptote towards the desired state.

For infrequent events such as large displacements in the low prior condition, the weight changes between the event and the output early in training are relatively large (left side of Figure 8c, inset). Taking a snapshot of performance at this stage would thus result in an apparent overweighting of their contribution to the output. This rule, however, predicts that once weights asymptote towards the desired state late in training (right side of Figure 8c, inset), events should contribute to the model’s performance in a manner proportional to their relative frequencies, and psychometric curves should become parallel to one another.

##### Simulations

We evaluated the model’s ability to explain the results by simulating its performance under experimental conditions. 95% confidence intervals for each simulated estimate were obtained by running 10000 simulations and identifying the middle 95% of each estimate, i.e., 2.5 percentile – 97.5 percentile. We simulated early prior training by following the same experimental structure as for the human experiments: a baseline block at *P*(*J*)=0.5 followed by two 600-trial blocks at *P*(*J*)=0.8 and *P*(*J*)=0.2, respectively. Of those trials, 70% were prior training trials at the lowest noise level, σ_target_=0.1°. The remaining 30% were medium (σ_target_=1°) and high-noise (σ_target_=2°) testing trials with a neutral movement statistic of 0.5 but simulated using the same inputs as the prior condition. Displacements were drawn from overlapping Gaussian distributions as in the experiments. Jumps were drawn from a distribution with μ_jump_= 0°, σ_jump_=2.5°, and non-jumps were drawn from a distribution with μ_non-jump_=0°, σ_non-jump_=0.5°. On trials where the target jumped, the desired state was set to 1 for the “jump” output unit and 0 for the “no jump” output unit. On trials where the target did not jump, it was set to 0 for the “jump” unit and 1 for the “no jump” unit. The learning rate was set at 0.5. Late training was simulated by allowing the model to run for 5000 trials and evaluating trials 3000-5000.

##### Combined (Discriminative + Bayesian ideal observer) Model Simulation

Finally, to better match observed patterns in the empirical data, we simulated a combined model whose final output was a weighted combination of outputs from the discriminative model, which incorporated visual noise, and a Bayesian ideal observer model which incorporated motor-driven noise. Motor noise was simulated in the Bayesian model by setting the width of the non-jump distribution, σ_non-jump_=1.5°, i.e., triple the width of the simulated experimental distribution to mimic saccadic suppression. Bayesian and discriminative model outputs were combined linearly, such that:

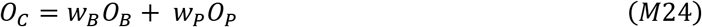

where *O_C_* is the output of the combined model, *O_B_* is the output of the Bayesian model, *O_P_* is the output of the discriminative model, and *w_B_* and *w_P_* are the weights assigned to the Bayesian and discriminative model, respectively. Further, the weights of the two component models added up to 1:

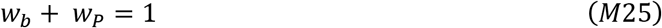

We combined the outputs of the Bayesian and discriminative models at relative weights of 0.1 and 0.9, respectively, to generate the data in Figure 8e.

## Supplementary figures

**Figure S1.**
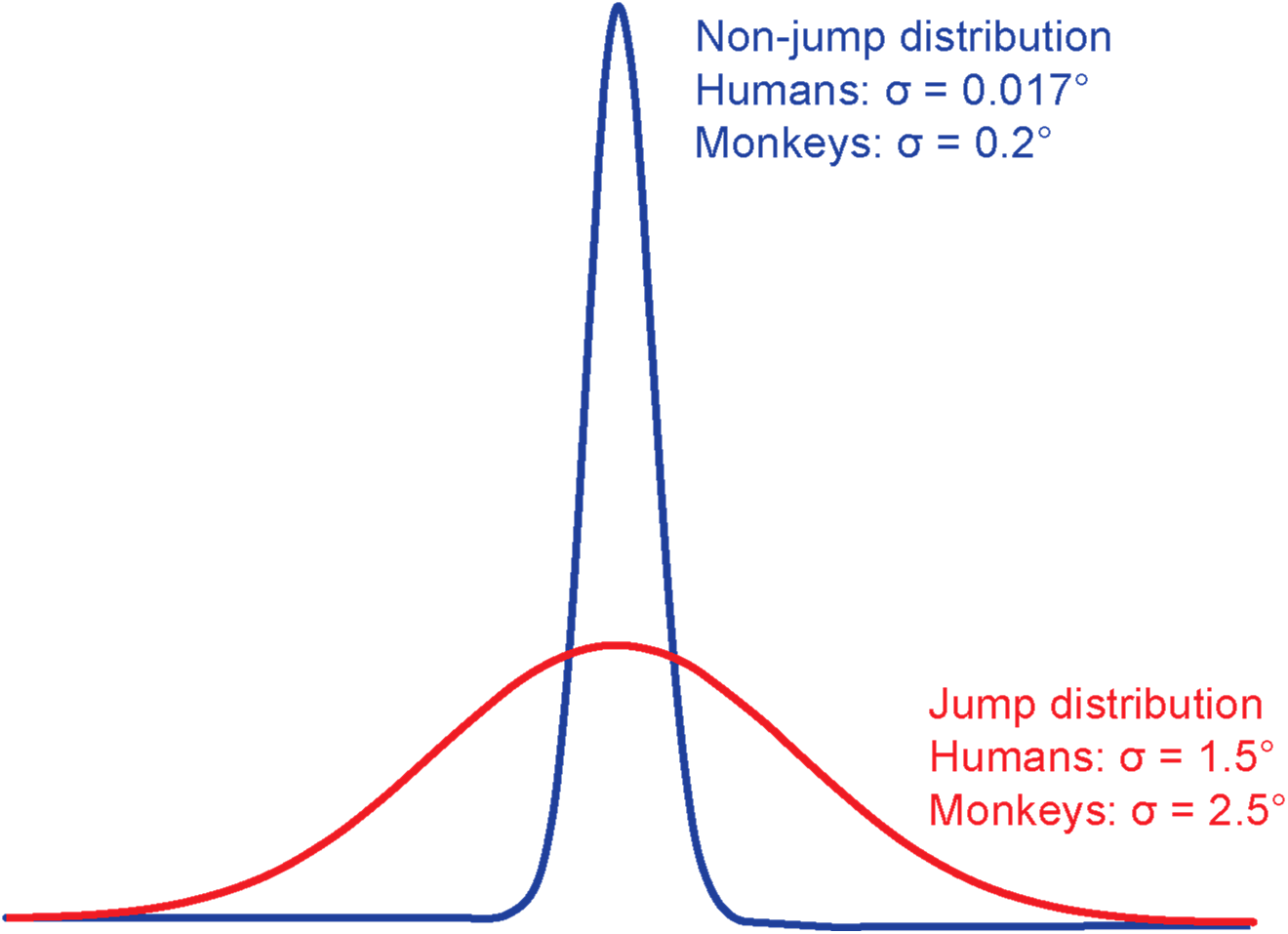
Schematized “jump” and “non-jump” distributions for the categorical experiments in humans (Experiments 1 and 2) and monkeys (Experiments 4, 5, and the control experiment).

**Figure S2.**
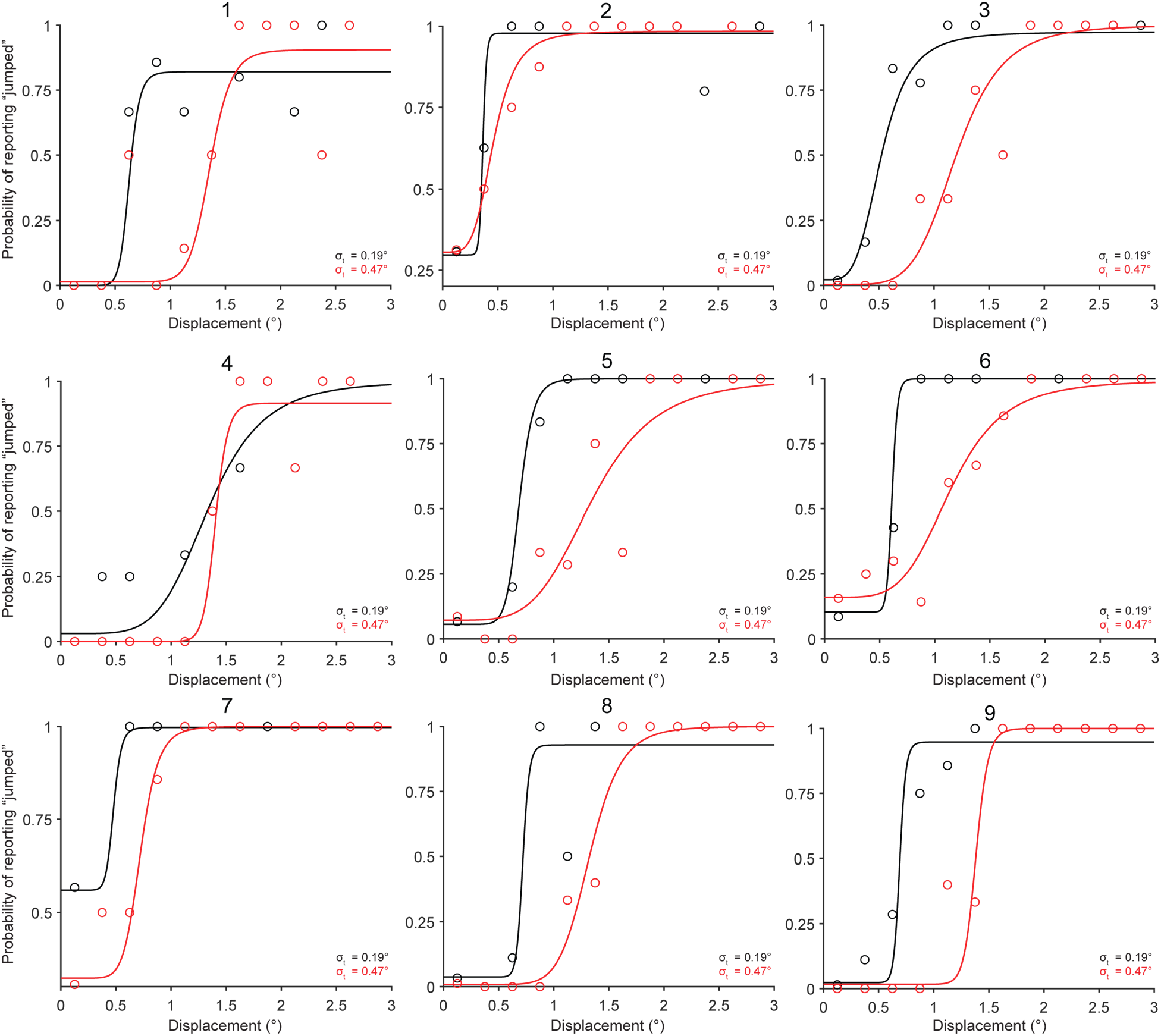
Individual participants’ psychometric curves for the Gaussian blob stimulus in Experiment 1.

**Figure S3.**
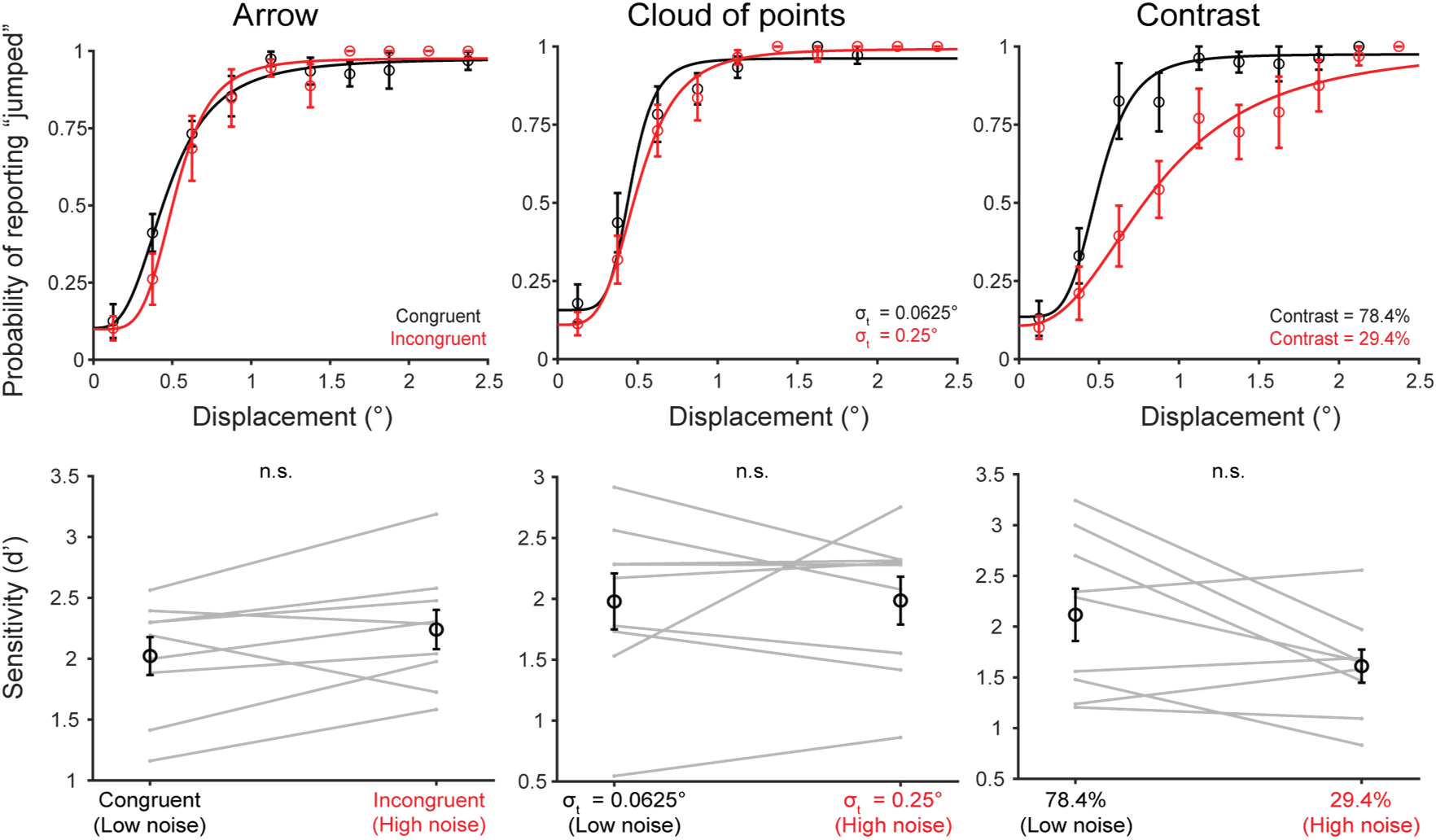
Data for the remaining stimuli tested in Experiment 1. Top row shows psychometric curves for each stimulus type and the bottom row shows d’.

**Figure S4.**
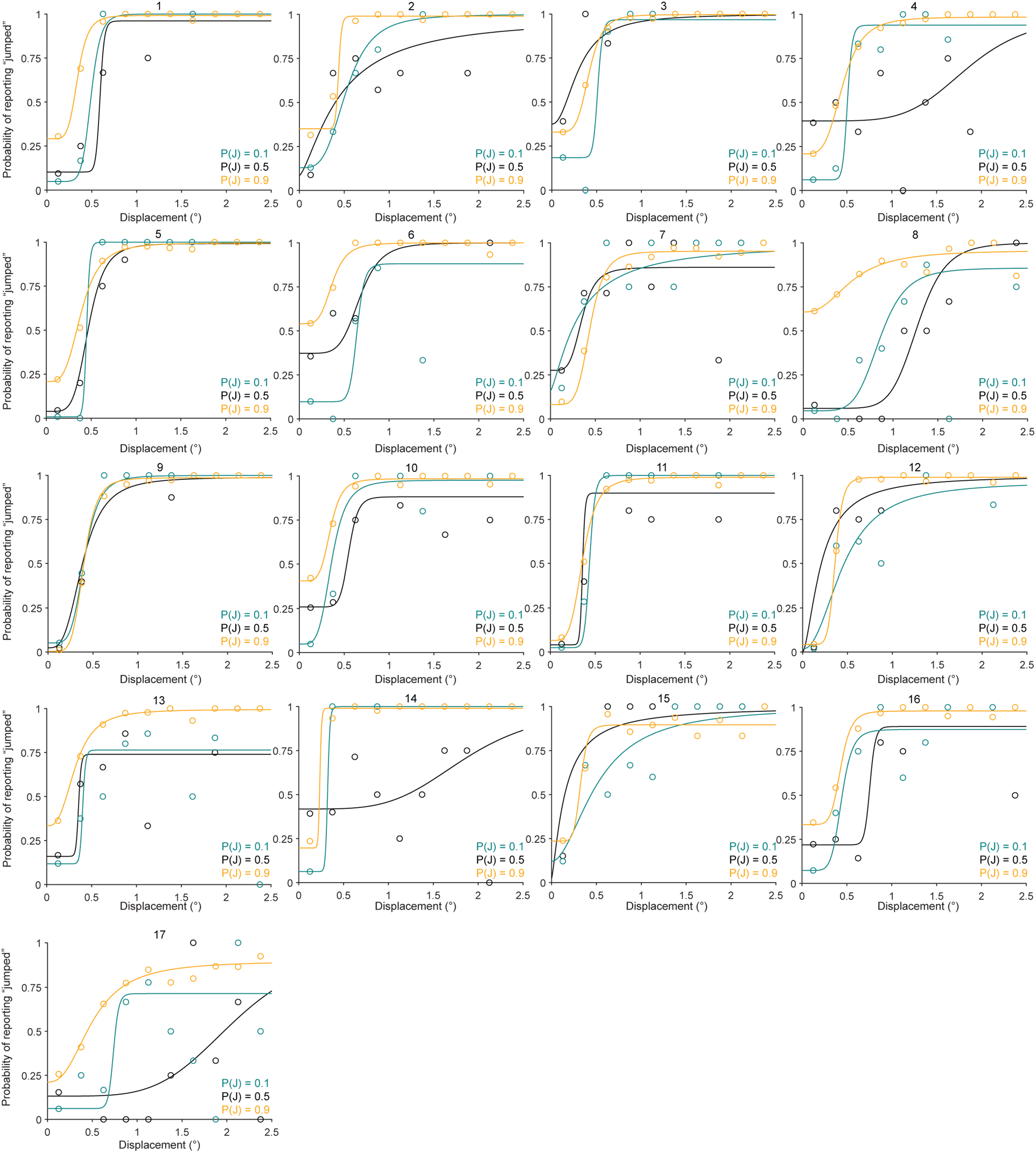
Individual participants’ psychometric curves for the prior training trials (low sensory noise) in Experiment 2.

**Figure S5.**
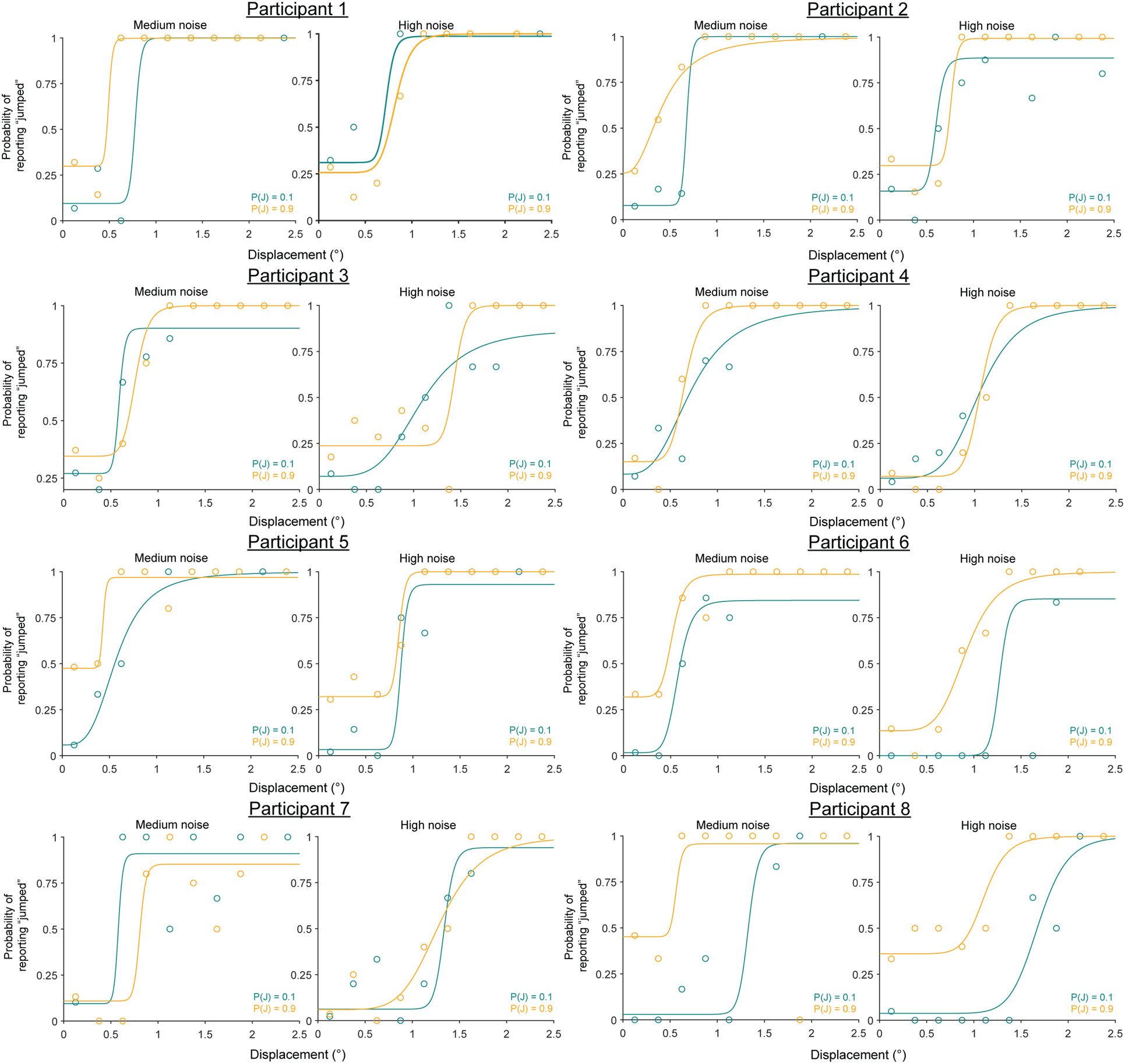
Individual psychometric curves for Participant 1 – 8 for the hypothesis testing trials with medium and high sensory noise in Experiment 2.

**Figure S6.**
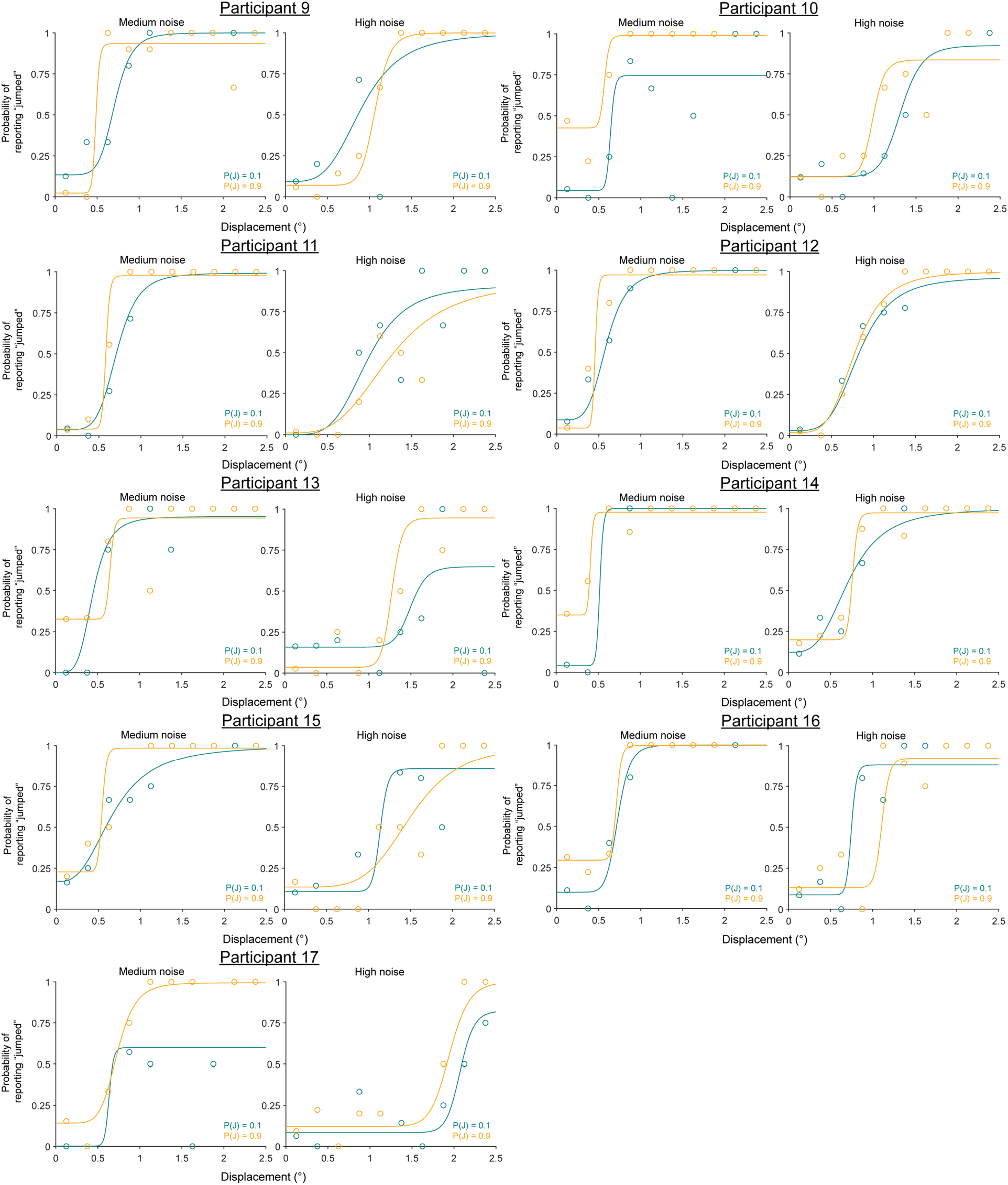
Individual psychometric curves for Participant 9 – 17 for the hypothesis testing trials with medium and high sensory noise in Experiment 2.

**Figure S7.**
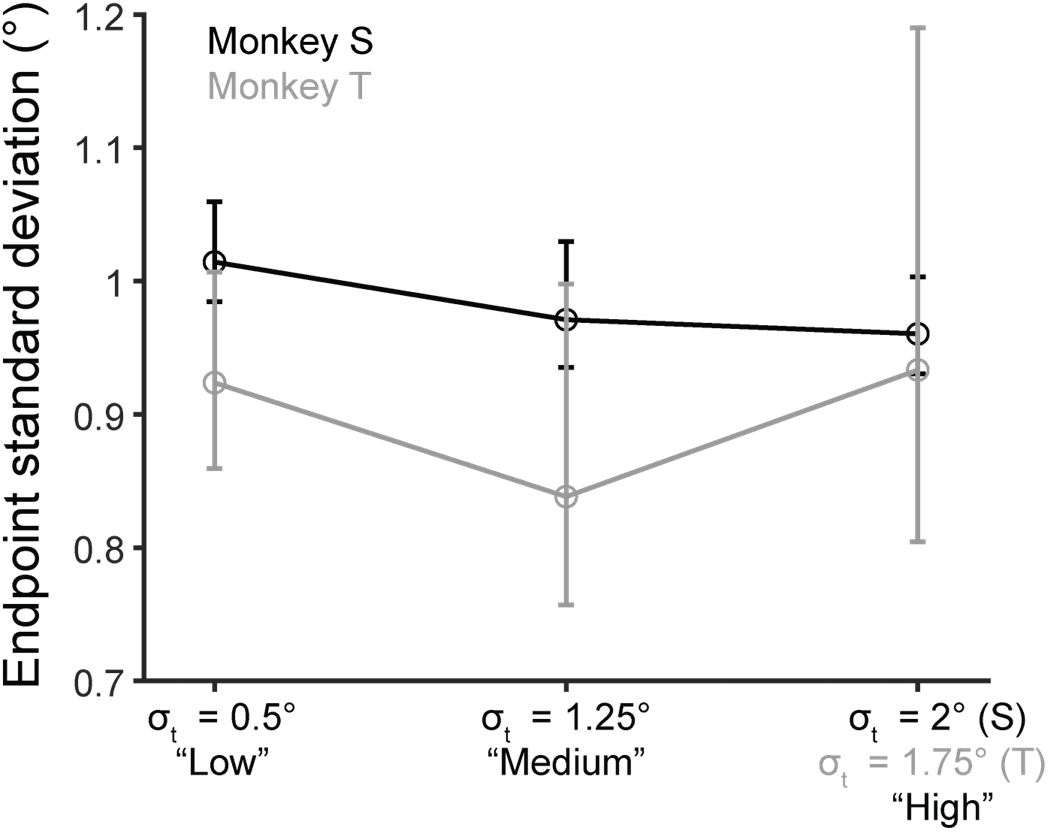
For both Monkey S (black)) and Monkey T (gray), 95 percent confidence intervals (CI) about endpoint standard deviations overlapped across noise levels. The standard deviations [CI] for Monkey S were 1.01 [0.98, 1.06], 0.97 [0.94, 1.03], and 0.96 [0.93, 1.00] for the low, medium, and high noise conditions. For Monkey T, they were 0.92 [0.86, 1.07], 0.84 [0.76, 1.00], and 0.93 [0.80, 1.19].

**Figure S8.**
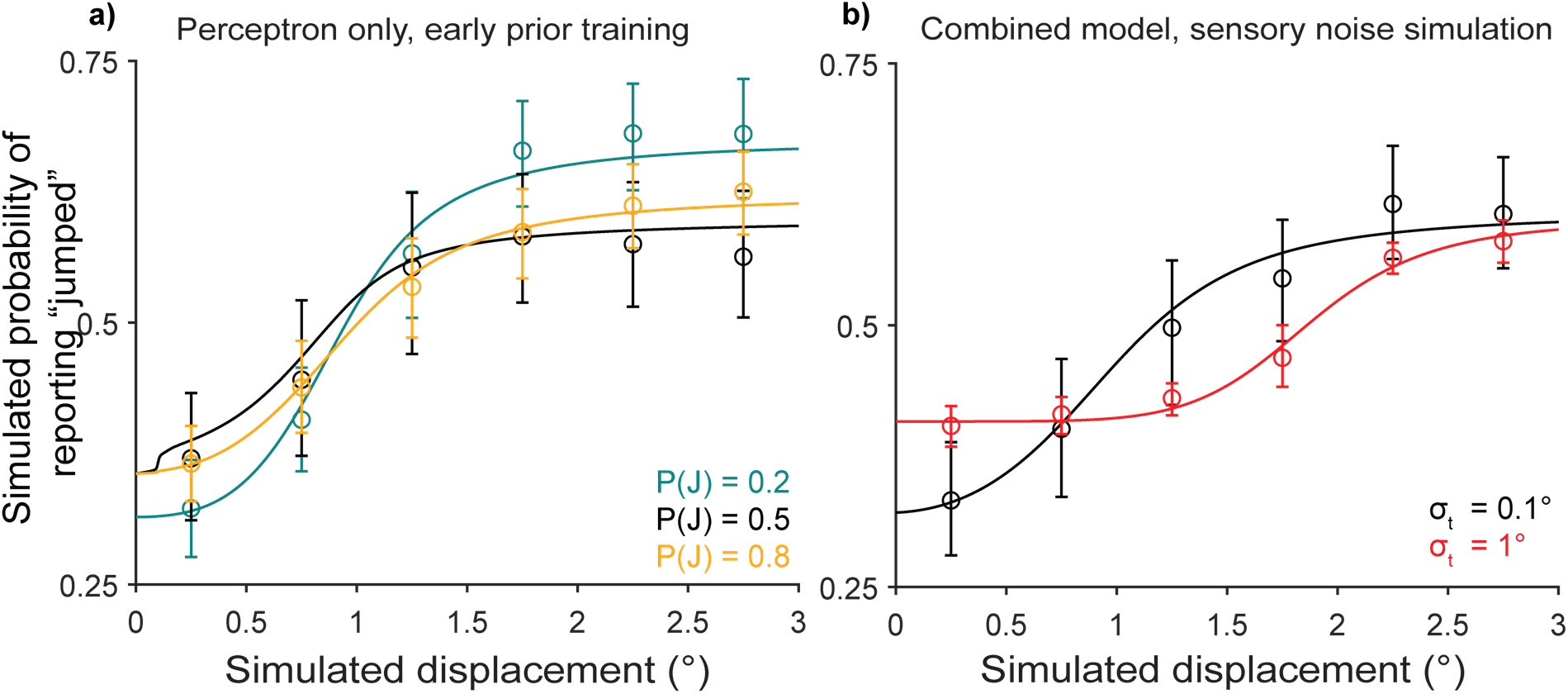
Simulations of early prior learning in the discriminative model alone (a) and two sensory noise levels in the combined (discriminative + Bayesian) model (b).

